# Genome sequence of 12 *Vigna* species as a knowledge base of stress tolerance and resistance

**DOI:** 10.1101/2022.03.28.486085

**Authors:** Ken Naito, Takanori Wakatake, Tomoko F. Shibata, Kohtaro Iseki, Shuji Shigenobu, Yu Takahashi, Eri Ogiso-Tanaka, Chiaki Muto, Kuniko Teruya, Akino Shiroma, Makiko Shimoji, Kazuhito Satou, Takashi Hirano, Atsushi J. Nagano, Norihiko Tomooka, Mitsuyasu Hasebe, Kenji Fukushima, Hiroaki Sakai

## Abstract

Harnessing plant genetic resources including wild plants enables exploitation of agronomically unfavorable lands to secure food in the future. The genus *Vigna*, family Fabaceae, consists of many species of such kind, as they are often adapted to harsh environments including marine beach, arid sandy soil, acidic soil, limestone karst and marshes. Here we report long-read assemblies of 12 *Vigna* species, achieving 95% or higher BUSCO scores. The comparative analyses discovered a new class of *WUSCHEL*-related homeobox (*WOX*) transcription factor superfamily that are incorporated into LTR retrotransposons and have dramatically amplified in some species of the genus *Vigna*. Except *WOX* transcription factors, however, gene contents are highly conserved among *Vigna* species with few copy number variations. On the other hand, transcriptome data provided some insights that transcriptional alterations played more important roles in evolution of stress tolerance in the genus *Vigna*. The whole genome sequences presented in this study will facilitate understanding genetic mechanisms of stress tolerance and application for developing new crops that are adapted to unfavorable environments.

## Introduction

To secure food for the global population, we have to recognize bottlenecks in agriculture. The first is that arable lands are only 10% of the global lands, and humans have cultivated almost all of it (Ritchie and Roser, 2013). The global lands are mostly covered with unfavorable soil, such as saline soil (~1.1 Gha) (Hassani et al., 2020), acidic soil (~4 Gha) (von Uexküll and Mutert, 1995) and alkaline calcareous soil (~3.5 Gha) (Hansen et al., 2006). In addition, 50% of the global lands are desert, where agriculture is impossible unless irrigated. However, irrigation brings other serious problems including further soil salinization (Hassani et al., 2021) and groundwater depletion (Pokhrel et al., 2021). Moreover, pests and diseases force farmers to lose 26% of their annual production (Cerda et al., 2017).

To overcome these bottlenecks, we have to harness the power of genetic resources including wild plants (McCouch et al., 2013). Modern cultivars have been intensely bred and thus are genetically vulnerable. In addition, modern agriculture largely relies on high-input farming with fertilizers, pesticides and other chemicals, which comes up with huge ecological costs beyond sustainability (Foley et al., 2011). Although crop yield per unit area has greatly increased in recent decades, we cannot expect the trend will continue in the next decade (Ray et al., 2012). As such, we have to exploit more unfavorable environments for agriculture in future. Solutions for an objective of such kind lie in wild plants that are naturally adapted to harsh environments.

As good examples of such wild species, we have been focusing on the genus *Vigna* (Tomooka et al., 2014). The genus consists of ~100 species, being a reservoir of diversity in a crop-rich family Fabaceae. Of them, not a few species are adapted to harsh environments including marine beach (Chankaew et al., 2014), arid sandy soil (Iseki et al., 2018), acidic soil (Tomooka et al., 2014), limestone karst (Takahashi et al., 2015) and marshes (Tomooka et al., 2014). In addition, resistance to various biotic stresses were also reported in some species (Birch Et al., 1986, Takahashi et al., 2019). As such, the genus *Vigna* could be a rich source of stress tolerance and resistance.

Several strategies are considered to harness the adaptability of the wild species. One is to introduce genes of stress tolerance into a crop species *via* cross breeding or genetic engineering. Simple crossing is still a powerful tool as many *Vigna* species are wild relatives of agronomically important crops such as cowpea (*Vigna unguiculata*), mungbean (*V. radiata*) and azuki bean (*V. angularis*) (Tomooka et al., 2014). Genetic engineering enables broader application of genes from wild species, although the target crop needs transformation techniques to be developed. The other is *de novo* domestication, which directly utilizes the naturally adapted wild species as a new crop. Instead of introducing stress tolerance from the wild to the domesticated, it needs to introduce domestication-related traits into the wild species (Takahashi et al., 2019). Given domestication-related traits have often arisen *via* loss-of-function mutations in single genes, *de novo* domestication is potentially a valuable option for developing a stress-adapted crop in a relatively short term (Takahashi et al., 2019).

To take any approaches described above, whole genome sequences are the most important basement to accelerate the processes. Although some domesticated species have already been sequenced (Lonardi et al., 2019, Knag et al., 2014, Sakai et al., 2015), no reference-level sequence is available on wild *Vigna* species except those we have sequenced previously (Takahashi et al. 2020, Takahashi et al. 2019). Thus, using PacBio long-reads, we sequenced and assembled the genomes of 9 more *Vigna* species (Summarized in Tables 1, S1, and Figs 1, S1). The assembled genomes achieved long contiguity, accurate gene annotation and enabled identification of syntenic blocks across all the species. We also identified private Pan-Genomic regions, which harbor ~1,000 genes per species. In addition, one gene family is highly expanded in a few species due to its incorporation into long terminal repeat (LTR) retrotransposons. We also performed transcriptome analyses and identified up-regulated genes that might have been involved in adaptation to harsh environments including beach and desert.

**Table 1.**
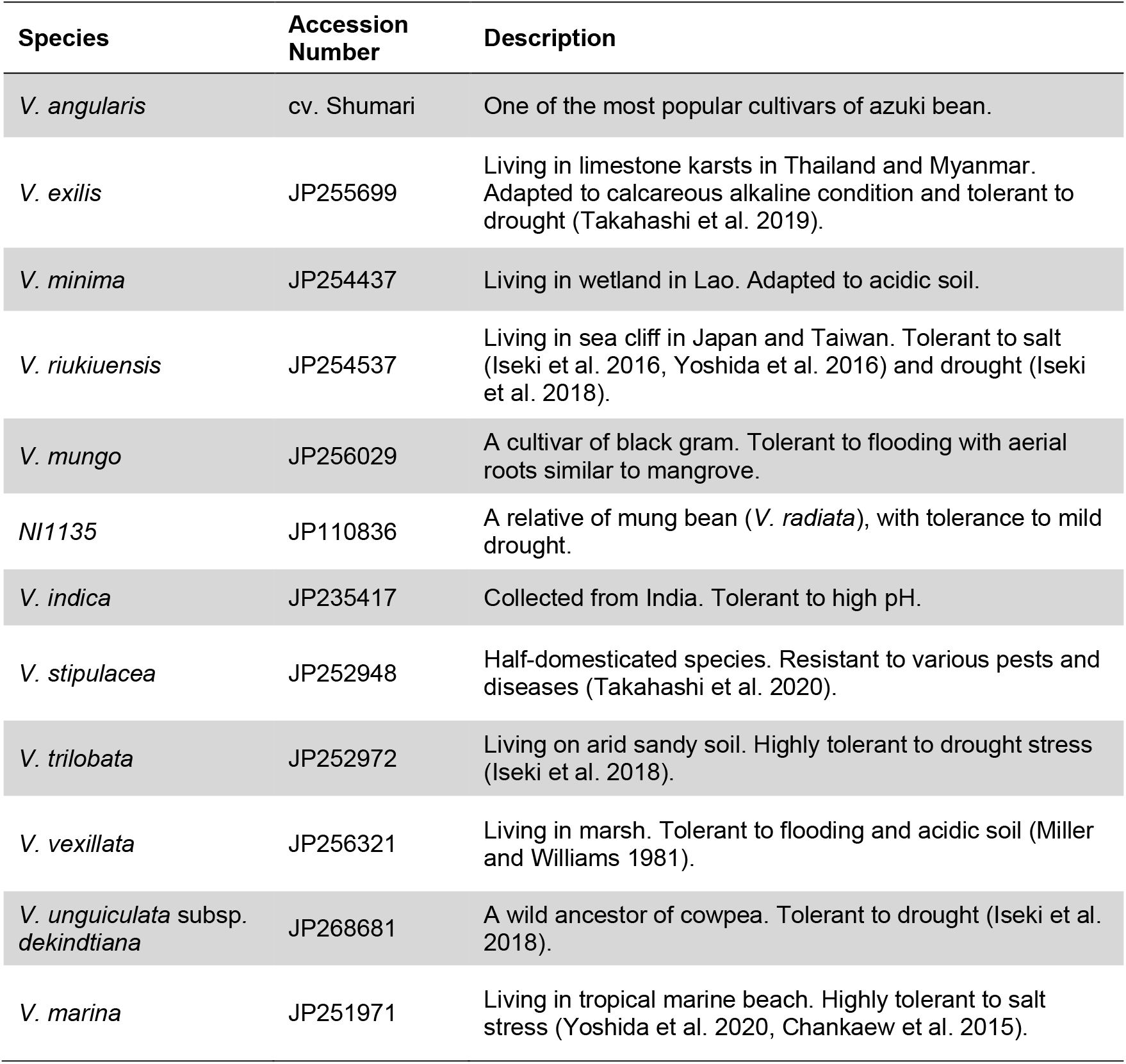
Plant materials.

**Figure 1.**
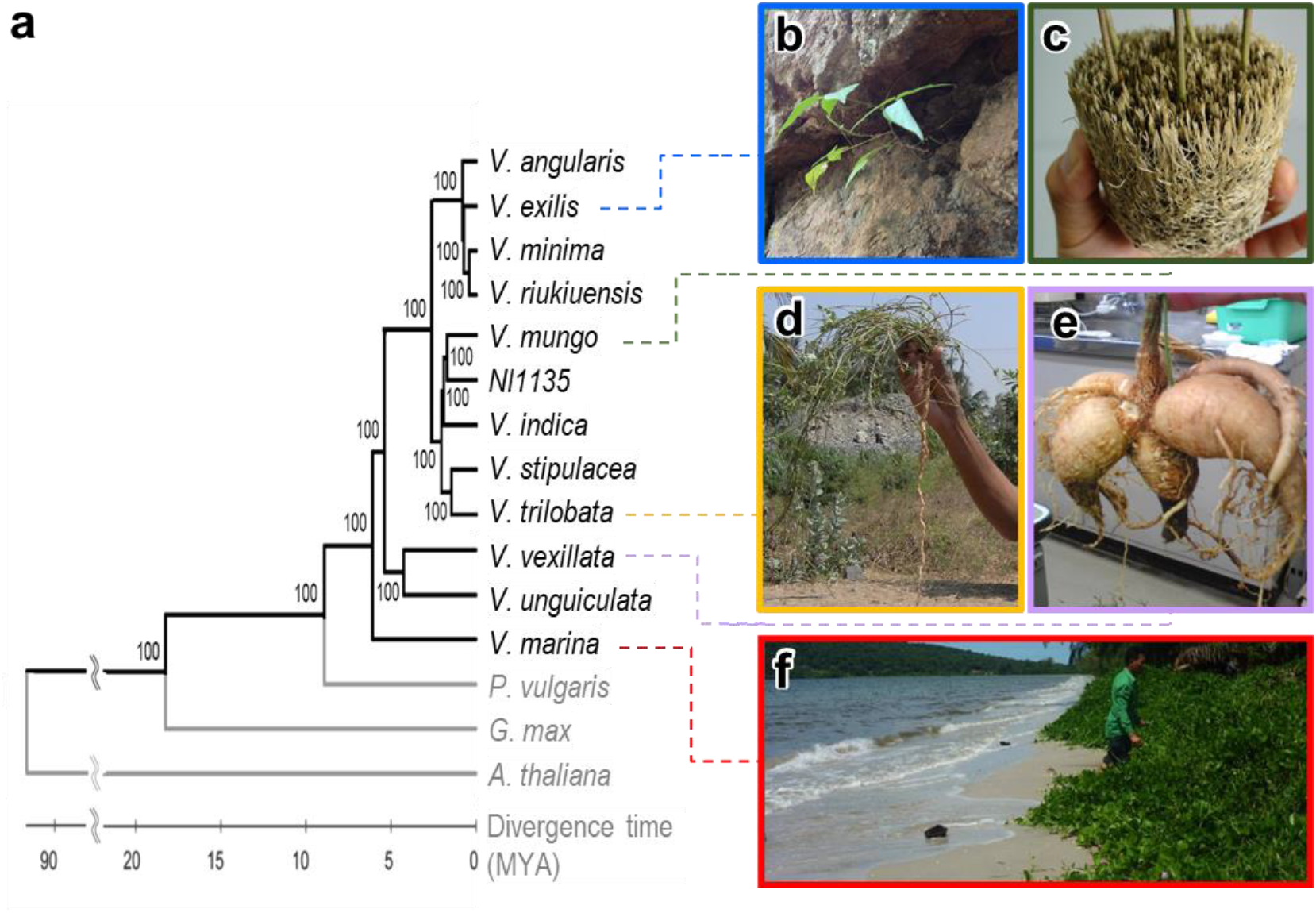
Phylogenetic tree based on the genomic data and photos of representative species. **a.** NJ tree of *Vigna, Phaseolus, Glycine* and *Arabidopsis*. Numbers beside branches represent bootstrap values (%) based on 1000 replications. **b.** *V. exilis* growing on a limestone rock. **c.** Root of *V. mungo* growing upward under flooded condition. **d.** Tap root of *V. trilobata* with few lateral roots. **e.** Tuber of *V. vexillata*. **f.** *V. marina* in a beach.

## Results

### De novo *assembly of 12 genomes*

In addition to our previous assemblies (Sakai et al. 2015, Takahashi et al. 2019, Takahashi et al. 2020), we sequenced and assembled the genome sequences of nine more species (Tables 2, S1). Of the 12, nine belonged to Asian Vigna (*Vigna angularis* (Willd.) Ohwi et Ohashi, *Vigna exilis* Tateishi et Maxted, *Vigna minima* (Roxb.) Ohwi et Ohashi, *Vigna riukiuensis* (Ohwi) Ohwi et Ohashi, *Vigna mungo* (L.) Hepper, *NI1135, Vigna indica* T.M. Dixit, K.V. Bhat et S.R. Yadav, *Vigna stipulacea* (Lam.) Kuntze and *Vigna trilobata* (L.) Verdc.), while three belonged to African Vigna (*Vigna vexillata* (L.) A. Rich., *Vigna unguiculata* subsp. *dekindtiana* (Harms) Verdc. and *Vigna marina* (Burm.) Merr.). Although the assemblies on African Vigna were relatively fragmented, those on Asians achieved higher contiguity and long terminal repeat assembly index (LAI) (Ou et al., 2018) (Table 2). Despite the difficulties in assembling the genomes of African Vigna, the annotated genes covered more than 94% of BUSCO genes in all the species except *V. unguiculata* ssp. *dekindtiana* (Table 2). In addition, gene contents and genomic syntenies were highly conserved among the species (Figs. S2, S3, Tables S2, S3). From the annotated genes of the 12 *Vigna* species, *Phaseolus vulgaris* L., *Glycine max* (L.) Merr. and *Arabidopsis thaliana* (L.) Heynh., we extracted 1,376 single copy orthologs and reconstructed a phylogenetic tree (Fig. 1). The result showed African Vigna shared basal lineages, whereas Asian *Vigna* have diverged more recently. It also showed *V. indica* formed a sister group with *V. mungo* and *NI1135*, although in our previous study it formed a sister group with *V. stipulacea* and *V. trilobata* (Takahashi et al. 2016) (Fig.1).

**Table 2.**
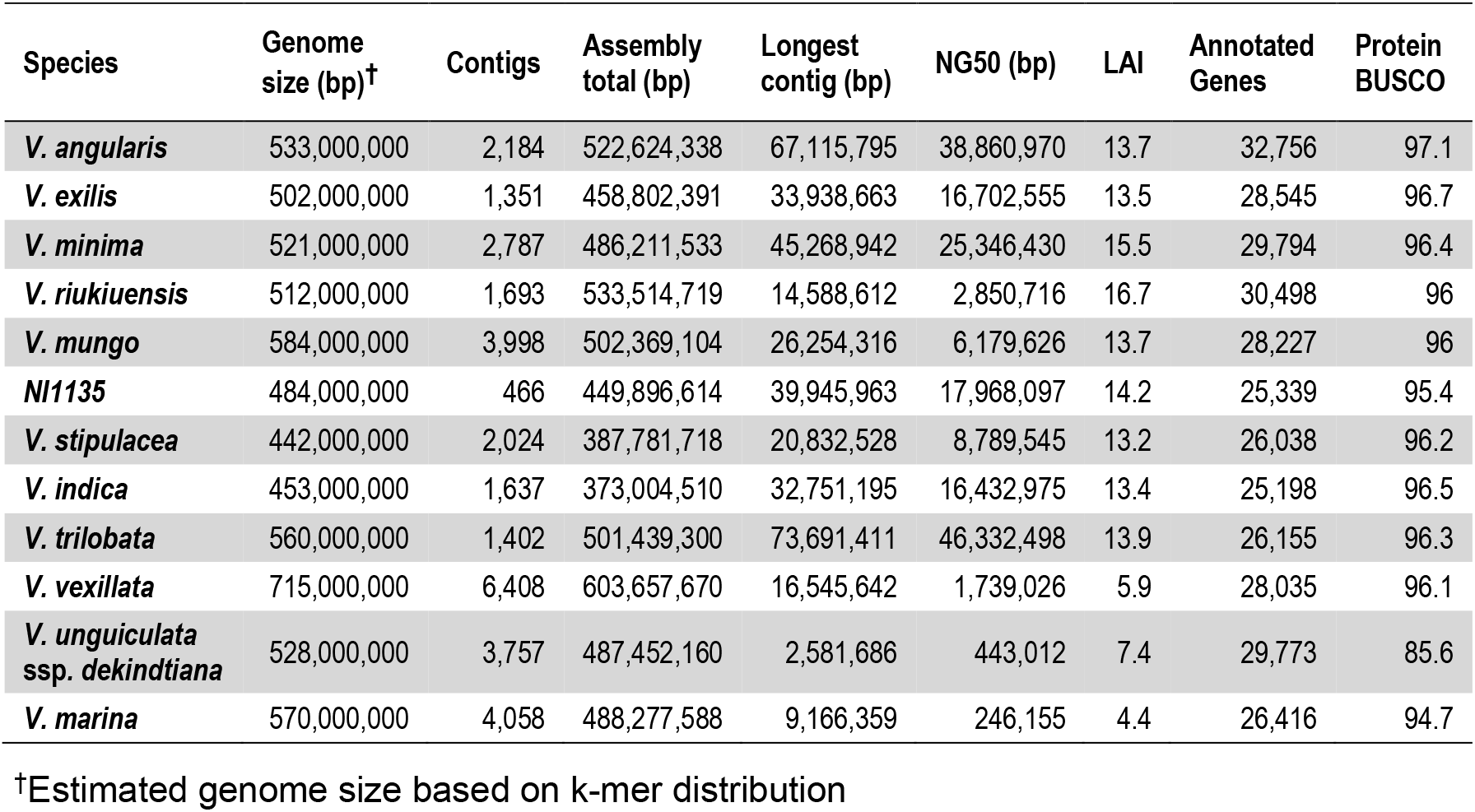
Stats of genome assemblies.

### Variations in TE abundance and insertions

We identified transposable elements (TEs) to assess their contributions to divergence of *Vigna* genomes. The results revealed TE contents basically correlated with genome size, ranging from 137 Mbp (37.0%) in *V. indica* to 302 Mbp (50.2%) in *V. vexillata* (Figs 2a, S4, Table S4).

**Figure 2.**
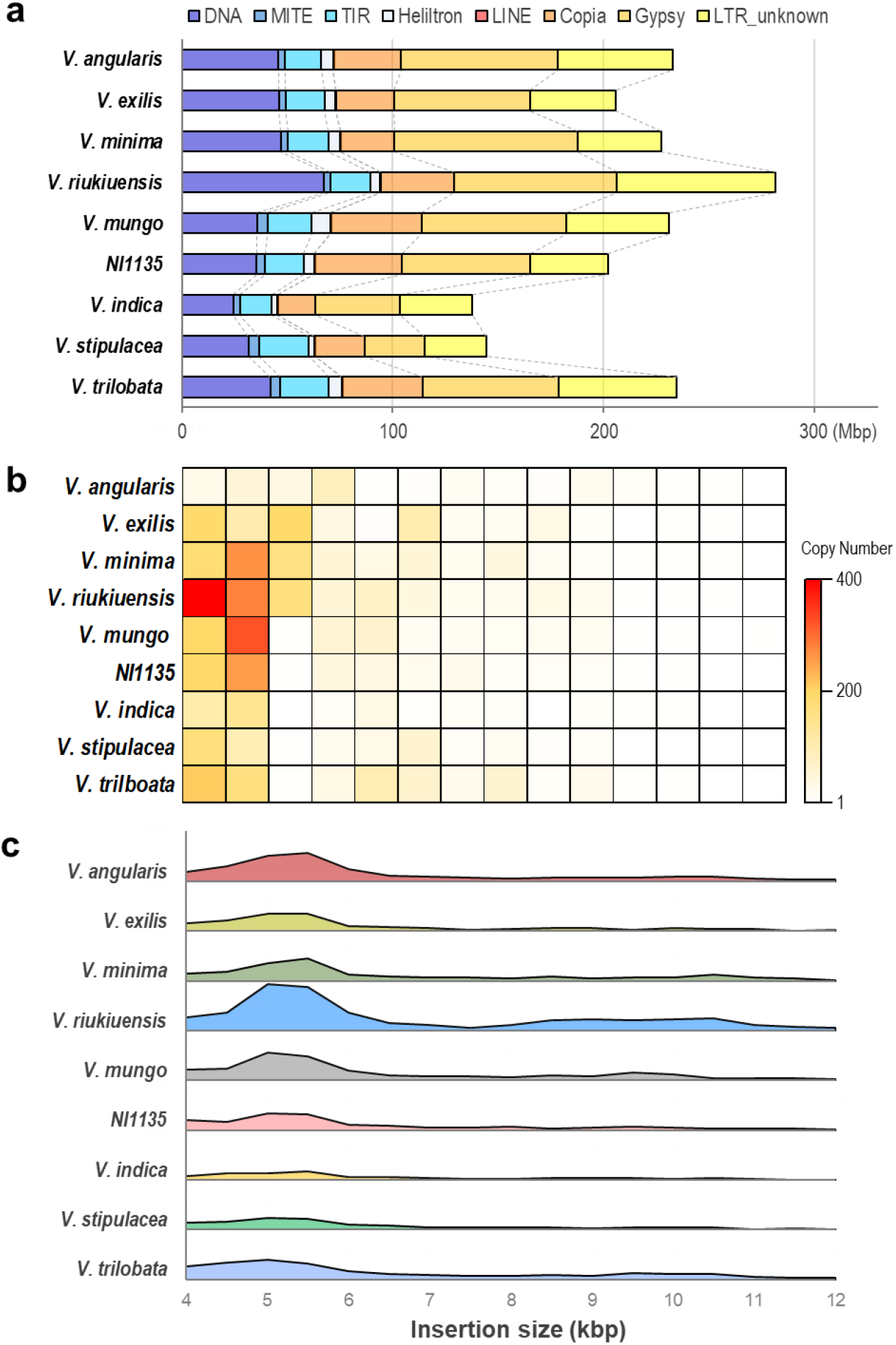
TE-related variations in Asian Vigna. **a.** TE contents in the genomes of 12 species. **b.** Copy number variations of TE-related orthogroups in Asian *Vigna* species. Each column indicates a TE-related orthogroup that are annotated as *GAG-POL-ENV* polyprotein or *TRANSPOSASE*. **c.** Presence variations (PVs) in Asian *Vigna* species. The range of Y-axes is 0-400.

In Asian Vigna, the TE abundance correlated with copy number of TE-related genes (transposases and gag/pol/env polyproteins) (Fig. 2b). However, few TE-related genes were annotated in the genomes of African Vigna, due to relatively fragmented assembly with lower LAI (Table 2).

Given there was a great variation in TE contents, we analyzed species-specific presence (insertion) variations (PVs) across the species. To do so, we created one-to-one alignment of each species pair and extracted unaligned fragments (≥500 bp) that were flanked with aligned sequences (≥5 kbp) in both ends without large gaps (≥100 bp). We excluded African Vigna from this analysis because we obtained few gapless alignments due to their diverged sequences.

The numbers of detected PVs basically correlated with TE abundance, being fewer (~2,000) in NI1135, *V. indica* and *V. stipulacea* and the most (4,206) in *V. riukiuensis* (Table S5). The size distributions of the PVs were overrepresented with ~5 kbp and ~10 kbp insertions in *V. riukiuensis*, suggesting potential amplification of TEs in these size ranges (Fig. 2c, Table S5).

### Pan-genomes and copy number variations highlighting TE hitchhiked by a host gene

The high coverage of annotated gene sets enabled us to perform a pan-genome analysis. As a first step, we identified private genes, which are present in only one species but not in others. The numbers of private genes varied among species, ranging from 668 in *V. indica* to 2,636 in *V. riukiuensis* (See the numbers of “species-specific” and “unassigned” genes in Tables S2, S3).

To presume functions of such private genes, we performed gene ontology (GO) enrichment analysis. Although few GO terms were enriched in those of most of the species, the GO terms “DNA-binding transcription factor activity” and “plant organ development” were highly enriched in *V. marina* and *V. riukiuensis* (Table S6). We extracted the private genes with these GO terms from the two species and found most of such genes were annotated as *“WUSCHEL-related homeobox* (*WOX*) transcription factors (Jha et al., 2020). In addition, 758 of the 2,636 private genes were such *WOX* transcription factors in *V. riukiuensis*, whereas 59 of the 1,649 private genes were so in *V. marina* (Fig. 3a).

**Figure 3.**
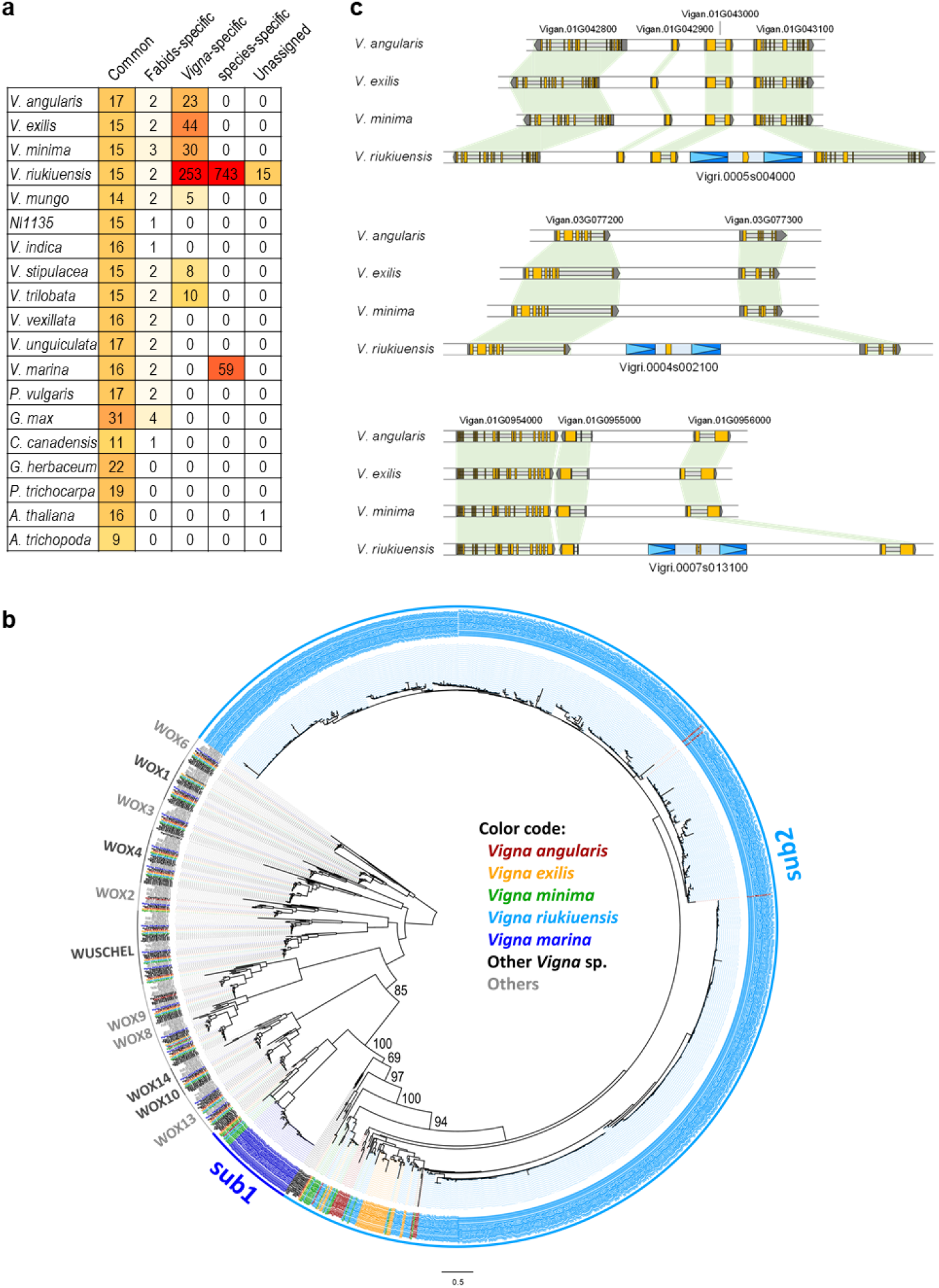
Amplification of WOX-related genes in the genus *Vigna*. **a.** Copy number variation in WOX-related orthogroups. b. A phylogenetic tree of *WOX* transcription factor superfamily. Bootstrap values are presented on Vïgna-specific clusters. Clustes of broadly conserved *WOX* genes are indicated with gene names in black or grey. Subclusters in V7gna-specific clusters are indicated as sub1 and sub2. c. Examples of genomic regions around sWOX gene loci in *V. riukiuensis* and the syntenic regions in other related species. Yellow and gray boxes indicate CDS and UTRs, respectively. Blue boxes with sky blue triangles indicate LTRs. Green shade indicates orthologous genes across species.

Thus, we extracted all the *WOX*-related genes including non-private ones (Fig 3a). As demonstrated in the gene tree in Fig. 3b, the “common” orthologous *WOX* genes were assigned to 9 clusters that consists of 1 or 2 genes from each species, whereas *Vigna-*specific *WOX* formed a huge cluster. This *Vigna*-specific cluster could be divided into two subclusters, one mainly with *V. marina* (sub1) and the other with *V. riukiuensis* (sub2). We note here that the both subclusters contained orthologs of most Asian Vigna in addition to those of *V. riukiuensis* or *V. marina* (Fig. 3b). Moreover, the gene tree revealed several events of dramatic amplification of these *Vigna-specific WOX* transcription factors. (Fig. 3b).

However, these *WOX* transcription factors were mostly single exon and encoded proteins of ~150 aa long. In addition, almost all the copies were not transcribed except a few, at least in our RNA-seq data. This contrasted with typical *WOX* transcription factors, which *are* often multi-exonic and encode proteins of 200-350 aa long. Thus, we designated the *Vigna*-specific WOX transcription factors as “*short WOX* (*sWOX*)”.

The dramatic amplification of *sWOX* genes in *V. riukiuensis*, together with abundance of PVs (Fig. 2c), intrigued us to test whether the *sWOX* amplification was related to PVs. As expected, at least 151 *sWOX* genes were located within the detected PVs in *V. riukiuensis* (Table S7). In addition, such *sWOX*-containing PVs were much larger (mainly 9-15 kbp) than the ORFs of *sWOX*. As the size of the PVs were similar to many typical retrotransposons, we further surveyed whether the *sWOX* amplifications were accompanied by any TE amplifications (Fig. 2a, Table S5).

As a result, we found in *V. riukiuensis* that 500 out of the 1,000 sWOX genes were embedded within intact LTR retrotransposons, which conserved not only complete pairs of LTRs but also target site duplications (TSDs) (Figs. 3c, S5). Moreover, syntenic relationships of the neighboring genes were highly conserved between *V. riukiuensis* and other related species (Fig. 3c). As such, the *sWOX* had been somehow inserted into LTR retrotransposons and then amplified through the copy-and-paste mechanism of retrotransposons.

However, it was not easy to find common features in the *sWOX*-harboring LTR retrotransposons. Although they were similar to each other in total length, there were great variations in the size and nucleotide sequence of the LTRs and TSDs (Fig. S5), despite the CDS of *sWOX* genes being highly conserved.

### Genes under positive selection

We also exploited *Vigna* genomes to identify genes under positive selection to understand the genetic basis underlying adaptability of *Vigna* species to harsh environments. To do so, we downloaded protein-coding sequences of 12 legume species (see methods). We ran OrthoFinder on the protein-coding sequences of the 24 species and extracted orthogroups that were aligned to each other without gaps of no more than 70 bp. We then performed HYPHY (Pond et al. 2005) and codeml (Yang 2007) to identify genes under positive selection specifically in the *Vigna* species.

As a result, we identified 34 genes that were positively selected (FDR < 0.05) in single species (Table S8). Of them, some are related to stress tolerance including *Radiation 51* (*RAD51*) (Doutriaux et al., 1998) and *RAD5* (Davies et al., 1994) in *V. mungo, Damaged DNA Binding 2* (*DDB2*) (Molinier et al., 2008) in *V. indica, Topoisomerase II (TOPII)* (Xie and Lam, 1994) in *V. marina*, and *N-Acetylglucosaminyl Transferase II (GNT-II)* (Yoo et al., 2021), in *V. trilobata*.

### Expression analysis of stress-related genes

Though we expected *Vigna* species had acquired stress tolerance by excess duplication of stress-related genes, we did not find any clear evidence of such events (Fig. S6). Thus, we suspected differences in gene expression profile could have played more important roles in evolution of stress tolerance in the genus *Vigna*. Although we had RNA-seq data of only non-stressed plants, we considered it was still possible to identify stress-related genes that were upregulated in the tolerant species. Thus, we assigned differentially expressed genes (DEGs) into 10×10 clusters by SOM clustering on the leaf expression dataset (Fig. S7) and the root expression dataset (Fig. S8).

To find any important genes in the tolerant species, we selected the well-characterized stress-related genes (Fig. 4, Table S9) from the clusters where gene expressions were higher in either of the species with salt tolerance (*V. riukiuensis* and *V. marina*) or drought tolerance (*V. exilis, V. riukiuensis, V. trilobata* and *V. unguiculata* ssp. *dekindtiana*). We do not mention other stresses here because we did not have systematic evidence regarding which species are tolerant or susceptible.

**Figure 4.**
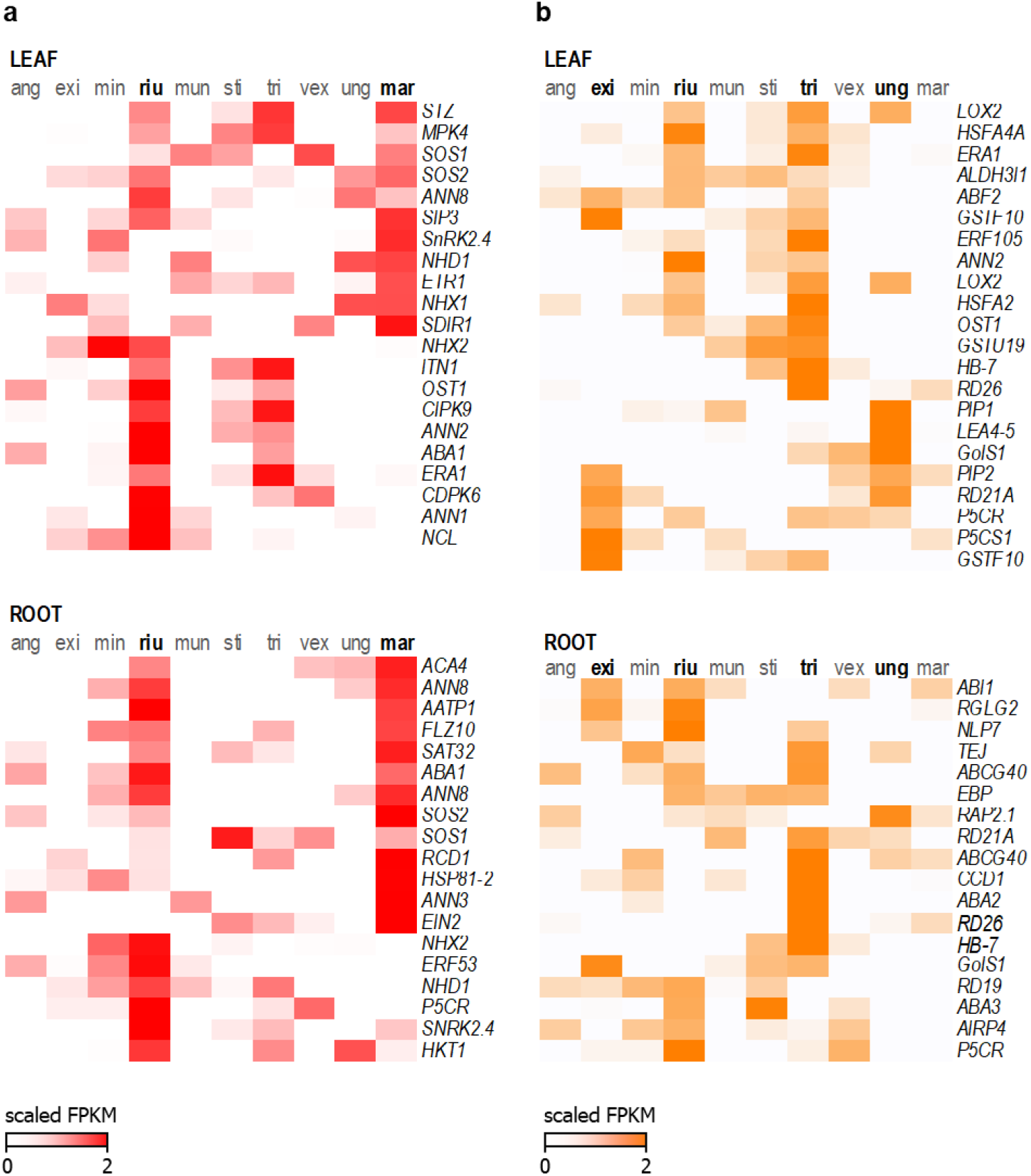
Expression of stress-related genes in *Vigna* species. a. Gens related to salt tolerance, b. Genes related to drought tolerance. The normalized FPKM values were log2-transformed and then centered with mean among species. Color scales indicate expression levels, where zero is the means. ang, exi, min, riu, mun, sti, tri, vex, ung and mar indicate *V. angularis, V. exilis, V. minima, V. riukiuensis, V. mungo, V. stipulacea, V. trilobata, V. vexillata, V. unguiculata* ssp. *dekindtiana* and *V. marina*, respectively.

As to salt tolerance, both the salt-tolerant species showed active transcription of several genes including the sodium transporter *Salt Overly Sensitive 1 (SOS1)* (Shi et al., 2000) and its activator *SOS2* (Liu et al., 2000) in both the leaf and the root (Fig. 4). In addition, *V. marina* actively transcribed additional sodium transporters such as *Na^+^/H^+^ Exchanger 1* (*NHX1*) (Apse et al., 1999) and *Sodium:Hydrogen Antiporter* (*NHD1*) (Müller et al., 2014) in the leaf, whereas *V. riukiuensis* actively transcribed potassium transporters such as *NHX2* (Cellier et al., 2004) and *High Affinity K^+^ Transporter 1 (HKT1)* (Sunarpi et al., 2005) in the root (Fig. 4). Genes related to ABA biosynthesis and signaling were also actively transcribed in *V. marina*, whereas those related to proline biosynthesis were actively transcribed in *V. riukiuensis* (Fig. 4, Table S9).

In drought-tolerant species, genes involved in ABA biosynthesis, transport and signaling, water transport, reactive oxygen species (ROS) scavenging and proline biosynthesis were actively transcribed in drought-tolerant species (Fig. 4, Table S9). Especially in the most drought-tolerant species *V. trilobata, Carotenoid Cleavage Dioxygenase 1* (*CCD1*) (Qin and Zeevaart, 1999) and *ATP-Binding Cassette G40* (*ABCG40*) (Kuromori and Shinozaki, 2010), which encode a key enzyme of ABA biosynthesis and an ABA transporter, respectively, were highly transcribed (Fig. 4). *V. riukiuensis* shared many of the upregulated genes in common with *V. trilobata*, whereas genes related to water transport and ROS scavenging were more transcribed in *V. unguiculata* ssp. *dekindtiana* and those related to proline biosynthesis were so in *V. exilis* (Fig. 4, Table S9).

## Discussion

In this study, we established a genomic basement of the genus *Vigna*, an important legume group as a genetic resource of stress tolerance. Thanks to the long-read sequencing, our assemblies achieved high contiguity, which enabled us to capture not only protein-coding sequences but also highly repetitive elements (Tables 2, S4). Now all the sequence data, including the gene annotation, are available *via* the *Vigna* Genome Server (*Vig*GS: https://viggs.dna.affrc.go.jp/) (Sakai et al. 2016).

The high-quality annotation enabled us to find *sWOX*, which is a new, *Vigna*-specific member of the *WOX* transcription factor superfamily. The *sWOX* gene copies mostly lack introns, suggesting it has originated *via* retroposition. The most interesting feature of *sWOX* is that it has been incorporated into LTR retrotransposons to amplify its copies in several species, especially in *V. riukiuensis* (Fig. 3). The phenomenon of host gene duplication *via* TE amplification system reminds us of *PACK-Mutator like Elements* (*PACK-MULEs*), a DNA transposon which has captured host gene sequences from multiple loci (Jiang et al. 2004). Our finding has added a new example of TEs (first as a retrotransposon) that have directly impacted gene contents in the host genome.

The amplification of *sWOX* in *V. riukiuensis* and *V. marina* (Fig 3a, c) brings an interesting question; does it have any relation with adaptation to coastal environments including high salinity? Though we do not currently have any direct evidence, several recent studies have reported some of the *WOX* transcription factors can confer osmotic tolerance by increasing root length (Wang et al., 2019) and root hair density (Cheng et al., 2016). In addition, as the *sWOX* genes have homeobox domain, it may bind to multiple loci and affect regulation of nearby genes when expressed. Thus, it would be worth testing whether overexpressing or knocking down *sWOX* genes affect root phenotypes and salt tolerance. On the other hand, it is also possible that the LTR retrotransposons harboring *sWOX* are simply activated by salt stress or other environmental cues in tropical coasts. In any case, further studies are necessary to test these hypotheses.

However, except *sWOX*, we identified few copy number variations especially in stress-related genes among *Vigna* species (Fig. S6). The result is not what we had expected before this study, as extremotolerant organisms including sleeping chironomid (Gusev et al., 2014) and tardigrade (Hashimoto et al., 2016) have expanded gene families of protectant proteins and ROS scavenging proteins. Similar cases have also been reported from plants, including *SOS1* expansion up to 14 copies in the coconut genome (Xiao et al., 2017). On the contrary, the conserved copy numbers of such genes across *Vigna* species suggests changes in transcriptional regulation have played more direct roles in acquiring stress tolerance, although we cannot rule out the possibility that such regulatory changes have arisen from expansion of transcription factors including *sWOX*.

Our expression data supported the above idea that evolution of gene regulation is important in stress tolerance of *Vigna* species (Fig. 4). Although our RNA-seq samples were limited to a non-stressed growth condition, many well-known genes involved in salinity tolerance or drought tolerance are upregulated in the salt-tolerant or drought-tolerant species, respectively (Fig. 4). Although some of the upregulated genes, such as *SOS1* and *SOS2*, are shared across the tolerant species, many others are specific to only one or a subset of the tolerant species (Fig. 4). This supports our conclusions in our preceding studies that tolerance to salinity and drought have independently evolved during diversification and speciation in the genus *Vigna* (Iseki et al., 2016, 2018). In any case, transcriptional alteration provides relatively a straight-forward explanation for the stress tolerance in the genus *Vigna*.

As demonstrated above, we consider wild plants including *Vigna* species to be useful materials to learn genetics of stress tolerance. Now scientists have identified thousands of stress-responsive genes in model plants, but the biggest problem is that little is known about how to use these genes to develop a practical stress-tolerant crop. To this end, we have to know which genes and how many of the thousands to be introduced to the target crop, or which tissues or which time the genes to be expressed. Our results provided some insight regarding which genes are important for adaptation to saline or arid environments (Fig. 4). Though we definitely need to elucidate regulation of gene expressions in higher resolution, it is possible because time-course sampling and single-cell RNA-seq are already feasible approaches with currently available techniques. Thus, the genome sequences we have assembled will be an important knowledge base for elucidating adaptation to agronomically unfavorable environments.

Last but not least, the genome sequences here will accelerate *do novo* domestication of wild *Vigna* species. It enables not only reverse genetic approaches to screen for mutants of known domestication-related genes, but identification of yet unknown genes by screening those that are degraded in domesticated species but conserved in the wilds. Or, as Takahashi et al. (2019) has demonstrated, it helps expand a catalog of domestication-related genes by identifying a responsible gene from a mutant that was screened for domestication-related traits in a forward genetic approach. Thus, our genome sequences will be a useful tool for developing new crops out of wild plants that are already well-adapted to unfavorable environments.

In this study we have sequenced only 12 species out of >100 in the genus *Vigna*. The wild species other than the 12 also have valuable traits and characters including stress tolerance (Iseki et al., 2016, Iseki et al., 2018, Yoshida et al., 2016, Yoshida et al., 2020), vigorous growth (Takahashi et al., 2015) and symbiosis with various rhizobia that are also naturally adapted to harsh environments (Mortuza et al., 2020). Further sequencing more species of the genus *Vigna*, together with many other plant taxa (Leebens-Mack et al., 2019), will facilitate fully understanding of adaptation to any desired environments.

## Experimental procedures

### Plant materials

All the accessions tested in this study, which are provided by National Genebank Program in Research Center of Genetic Resources, NARO, are summarized in Table 1 and Fig. S1. We extracted genomic DNA with standard CTAB method and purified it with QIAGEN Genomic-tip 20/g (Qiagen KK, Tokyo, Japan).

We also extracted RNA for gene annotation and transcriptome analysis. For the former purpose, we sowed the seeds in pots filled with culture soil and grew the plants in a greenhouse. We sampled the whole shoots and the whole roots from the 3-week-old plants (no replicate). For the latter purpose, we grew plants with hydroponic culture in a growth chamber (14h light at 28°C and 10h dark at 24°C) and sampled leaves and roots from the 3-week-old plants (triplicates). We used RNeasy Plant Mini Kit (Qiagen KK, Tokyo, Japan) to extract RNA.

We also cultivated 190 F2 plants derived from two accessions of *V. trilobata* (JP210605 and JP252972) in a greenhouse, sampled primary leaf of each plant and extracted DNA with DNeasy Plant Mini Kit (Qiagen KK, Tokyo, Japan).

### DNA and RNA sequencing

The extracted DNA for genome assembly was sheared into 20 kb fragments using g-TUBE (Covaris, MA, USA) and converted into 20 kb SMRTbell template libraries. The library was size-selected for a lower cutoff of 7 kb using BluePippin (Sage Science, MA, USA). Sequencing was performed on the PacBio RS II using P6 polymerase binding and C4 sequencing kits with 180 min acquisition.

The DNA was also used for sequencing with Illumina HiSeq 2000 platform. Library construction and sequencing were provided as a custom service of Eurofins MWG GmbH (Ebersberg, Germany). Sequencing libraries included a paired-end library of 300 bp inserts. For *V. marina* and *V. trilobata*, mate-pair libraries of 3 kb, 8 kb and 20 kb were also constructed. One lane of the flow cell was used for each library.

The RNA was sequenced with DNBSEQ-G400RS platform. Library construction and sequencing were provided as a custom service of GeneBay Inc. (Yokohama, Japan).

The DNA of *V. trilobata’s* F2 plants were processed for RAD-seq. The DNA was double-digested with EcoRI and BglII (New England Biolabs, Ipswich, MA, USA), size-selected for 320 bp, pooled and pooled as described by Sakaguchi et al. (2015). The library was sequenced with an Illumina HiSeq 2000 sequencer (Illumina, San Diego, CA, USA) with 51-bp single-end reads.

### Genome assembly and scaffolding

The obtained reads were assembled with Celera Assembler 8.3rc1 (Berlin et al. 2015) with ‘asmOvlErrorRate = 0.1, asmUtgErrorRate = 0.06, asmCgwErrorRate = 0.1, asmCnsErrorRate = 0.1, asmObtErrorRate = 0.08, utgGraphErrorRate = 0.05, utgMergeErrorRate = 0.05’ options. For *V. trilobata* and *V. marina*, the assembled contigs were scaffolded using SGA ver. 0.10.1341 (Simpson and Durban 2012) on Illumina mate-pair reads. We ran PBJelly2 (English et al. 2012) three times to close as many sequence gaps as possible using the error-corrected PacBio reads. We then polished the assembled contigs and scaffolds with short reads using Pilon 1.20 (Walker et al. 2014).

For further scaffolding *V. trilobata*, we constructed a genetic map according to the methods described in Marubodee et al. (2015). We did RAD-seq and mapped the obtained sequences to the scaffolds of *V. trilobata* with bwa-0.712 (Li and Durbin 2009) and genotyped the F2 plants with stacks-1.48 (Catchen et al. 2013) with default settings. We then built a genetic map using AntMap (Iwata and Ninomiya, 2006) (Fig S9), and manually surveyed discordance between the assembly and the genetic map. We discarded the discordant regions as misassemby, and then anchored the corrected scaffolds/contigs onto the linkage map, as previously described by Sakai et al. (2015). For other species, we constructed the scaffolds by Reference-Assisted Chromosome Assembly (RACA) program v.0.9.1.1 (Kim et al. 2013) using the genome sequences of *V. angularis* and *P. vulgaris* as the reference and outgroup species, respectively.

### Annotation

We performed *ab initio* gene prediction with BRAKER version 1.6 (Hoff et al., 2016) with RNA-Seq data. Besides, we predicted gene structures by genome-guided and *de novo* assembly approaches using TopHat 2.1.0 (Kim et al., 2013), Cufflinks 2.2.1 (Trapnell et al., 2010), Trinity 2.1.1 (Grabherr et al., 2011), and PASA pipeline 2.0.2 (Haas et al., 2008). We used Transdecoder 2.0.1 (https://github.com/TransDecoder/TransDecoder) and Trinotate 2.0.2 (Bryant et al., 2017) to predict ORFs. We also did protein mapping approach using Exonerate 2.2.0 (Slater and Birney, 2005) to map protein sequences of the *Glycine max* (Wm82.a2.v1) (Valliyodan et al. 2019), *Phaseolus vulgaris* (v.1.0) (Schmutz et al. 2014), *Medicago truncatula* Gaertn. (Mt4.0v1) (Tang et al., 2014), and *V. angularis* (Willd.) Ohwi & H.Ohashi (VANGULARIS_V1.A1) (Sakai et al., 2015) to the genome assemblies. The protein sequences were downloaded from Phytozome (*G. max, P. vulgaris, and M. truncatula*) (Goodstein et al., 2012) and *VigGS* (*V. angularis*) (Sakai et al., 2016). We combined the *ab initio* gene models, transcript alignments and protein alignments by EvidenceModeler 1.1.1 (Haas et al., 2008) and updated the predicted gene models by PASA (Haas et al., 2003), with manual curation on gene models with extremely long introns and those merged by PASA. We used BUSCO v4 (Waterhouse et al., 2017) to evaluate protein sequences of annotated genes. Syntenic blocks were identified by MCScanX (Wang et al., 2012) and visualized by SynVisio (Bandi and Gutwin, 2020).

We also evaluated the genome assembly by Long Terminal Repeat Assembly Index (LAI) (Ou et al., 2018) and annotated transposable elements (TEs) by running The Extensive *de novo* TE Annotator (EDTA) with default settings (Ou et al. 2019).

### Species tree inference

In order to construct the phylogenetic tree of the 12 *Vigna* species and two legume species, *G. max* (Valliyodan et al. 2019) and *P. vulgaris* (Schmutz et al. 2014), we conducted Orthofinder 2.3.3 (Emms and Kelly, 2019) and identified single-copy orthogroups. *Arabidopsis thaliana* (Lamesch et al. 2012) was included in the analysis as an outgroup. For each single-copy orthogroup, we made a codon alignment using MAFFT 7.294 and removed any sites including gaps using Gblocks 0.91b (Castresana 2000). The trimmed alignments were converted into codon alignments by PAL2NAL (Suyama et al. 2006) and concatenated, which was used as an input for species tree inference. We adopted the modified Nei-Gojobori method (Zhang et al. 1998) to calculate the synonymous distance matrix and constructed the phylogenetic tree using the neighbor-joining method (Saitou and Nei 1987). Divergence time was estimated based on a rate estimate of 6.5 x 1.0^-9^ substitutions per site per year (Gaut et al. 1996).

### Detecting presence variations (PVs)

In order to detect species-specific PVs, first one-to-one genome alignment of each pair of species was created by LAST (Firth and Noé 2014). Second, for each pair of species, unaligned regions (≥500bp) in one species where flanking sequences (≥500bp for each end) were aligned with another species with no large gaps (>100bp) (PVs candidates) were extracted. Coordinates of the PVs were determined by aligning each PV and flanking sequences of the two species by MAFFT (Katoh and Standley 2013). Finally, species-specific PVs were defined as PVs verified among one or more species in both the same taxonomic section and other section.

### WOX *gene tree inference*

To extract all the WOX superfamily genes, TBLASTX search was performed against CDS sequences from 19 angiosperms (12 *Vigna* species, *P. vulgaris* (Schmutz et al. 2014), *G. max* (Valliyodan et al. 2019), *Cercis canadensis* L. (Griesmann et al. 2018), *Populus trichocarpa* Torr. & A.Gray ex Hook. (Tsukan et al. 2006), *Gossypium herbaceum* L. (Huang et al. 2020), *A. thaliana* (Lamesch et al. 2011), and *Amborella trichopoda* Baill. (Amborella Genome Project, 2013)) with e-value threshold = 0.01 and minimum coverage threshold = 0.25 using 16 *WOX* superfamily genes from *V. riukiuensis* and *V. marina* (6 conserved genes among angiosperms, 5 Fabaceae specific genes, 5 genes from *V. riukiuensis-specific* subfamily) as queries. BLAST hits (1,548 genes) were aligned in-frame using mafft 7.480 (Katoh and Standley, 2013) and tranalign in EMBOSS 6.6.0 (Rice et al. 2000). Sequences with many gaps were removed using MaxAlign 1.2 (Gouveia-Oliveira et al. 2007). Less-alignable codon sites were removed with ClipKIT 0.1.2 (Steenwyk et al. 2020) with the default parameters. The processed alignment was used as an input for the maximum likelihood tree reconstruction with IQ-TREE 2.1.2 under the GTR + G4 model (Nguyen e al. 2015). The obtained gene tree was reconciled with the species tree using GeneRax 2.0.2 (--rec-model “UndatedDL”, --strategy “SPR”, --per-family-rates) (Morel et al., 2020). The species tree was reconstructed with IQ-TREE 2.1.2 under the GTR+F+R6 model selected by the ModelFInder using shared single BUSCO genes among 19 species. The embryophyta_odb10 was used as the lineage dataset for BUSCO analysis (v5.1.2) (Manni et al., 2021). Shared single BUSCO genes (417 genes) were processed the same as above-mentioned WOX superfamily genes and used as an input for IQ-TREE. The constraint tree option was used to follow the phylogenetic framework of APG IV at the order level (The angiosperm phylogeny group, 2016). The obtained species tree had the same topology in the Vigna clade as Fig. 1.

### Detecting positively-selected genes

In order to search for species-specific genes that have experienced positive selection, we basically followed the method described in Jebb et al. (2020). First, we conducted OrthoFinder 2.4.0 (Emms and Kelly, 2019) for 24 legume species consisting of 12 *Vigna* species, *Arachis hypogaea* L. (Bertioli et al., 2019), *Cajanus cajan* (L.) Millsp. (Varshney et al., 2012), *Cercis canadensis* L. (Stai et al., 2019), *Chamaecrista fasciculata* (Michx.) Greene (Griesmann et al., 2018), *Cicer arientinum* L. kabuli (Varshney et al., 2013), *Glycine soja* Siebold & Zucc. (Valliyodan et al., 2019), *Lotus japonicus* (Regel) K.Larsen (Kamal et al., 2020), *Lupinus albus* L. (Hufnagel et al., 2020), *Medicago truncatula* Gaertn. (Tang et al., 2014), *P. vulgaris* (Schmutz et al., 2014), *Pisum sativum* L. (Kreplak et al., 2019), *and Trifolium pratense* L. (De Vega et al., 2015). Protein sequences of the species other than 12 *Vigna* species were obtained from Phytozome (Goodstein et al. 2012). Second, we selected the orthogroups that included only single-copy orthologs in 12 *Vigna* species and then reconstructed rooted maximum likelihood (ML) gene trees using MEGA X (Kumar et al. 2018) and Notung 2.9.1.5 (Chen et al., 2000). Third, we used HYPHY package 2.5.17 (Pond et al. 2005) with aBSREL model and detected the genes showing signatures of selection (false discovery rate (FDR) < 0.05). We further performed the branch-site test implemented in codeml of PAML package 4.9j (Yang 2007) and selected the candidate genes (FDR < 0.05). Finally, we manually checked the multiple sequence alignments of the candidate genes and discarded the alignments including large gaps around the predicted positively-selected sites.

### GO enrichment analysis

We conducted OrthoFinder 2.4.0 on 12 *Vigna* species as well as *A. thaliana, G. max, and P. vulgaris* and identified orthogroups present only in single *Vigna* species and unassigned genes as species-specific genes. Besides, we performed InterproScan 5.51-85.0 (Jones et al. 2014) to assign the Gene Ontology (GO) terms to all genes. For each GO term assigned to species specific genes, we counted the numbers of genes having the GO term and genes not having the GO term for both species-specific genes and other genes. Then we tested the difference in the gene numbers by Fisher’s exact test and adjusted the *p*-values by Bonferroni correction method.

### Gene expression analysis

RNA-seq reads were mapped to transcriptomes of individual species using kallisto 0.46.2. Read counts were then subjected to the cross-species normalization by trimmed mean of M-values (TMM) (Robinson and Oshlack 2010). The TMM normalization factors were calculated from the expression levels of 9,270 single-copy orthologs identified by OrthoFinder 2.5.2 (Emms and Kelly, 2019). The normalization factors showed a consistent trend not by species but by organs (leaf < 1 and root > 1), suggesting that our RNA-seq data are comparable across *Vigna* species. TMM-normalized counts were converted to fragments per kilobase million (FPKM) and then transformed to log(X+1) values. The normalized FPKM values were filtered by edgeR (Robinson et al. 2010) for differentially expressed genes, and then clustered into 100 clusters by self-organizing map (SOM) clustering (Wehrens and Buydens 2007).

## Acknowledgement

This study was supported by grants from the Project of the NARO Bio-oriented Technology Research Advancement Institution (Research program on development of innovative technology) (H.S.), NIBB Collaborative Research Program (H.S.), JST PRESTO (K.N.), MEXT/JSPS KAKENHI grant number 22128001 (M.H.), and Moonshot R&D Program for Agriculture, Forestry and Fisheries (K.N.). We are also grateful for Ms. Shoko Ohi for her sophisticated operation of PacBio sequencer.

## Author contributions

KN and HS conceived and supervised the study. SS, MH, KS and TH coordinated the sequencing with help from CM, KT, AS and MS. KN and HS performed assembly and annotation. KI and AJN constructed genetic maps. YT, EOT, CM, AJN and KF generated transcriptome data. KN identified syntenic relationships of each gene. HS identified presence variations and positively-selected genes. TW and KF performed phylogenetic analyses. KN and TW performed clustering of transcriptome data. KN wrote the manuscript with input from TW, KF and HS.

## Supporting Information

Figure S1. Introduction of the wild species sequenced in this study.

Figure S2. Orthologous gene families in *Vigna, Phaseolus, Glycine* and *Arabidopsis*

Figure S3. Synteny plot between *V. angularis* and *V. minima, V. trilobata* or *V. unguiculata*. Figure S4. TE contents in the genomes of 12 *Vigna* species.

Figure S5. Close-ups of the LTR retrotransposons and the harbored *sWOX* genes in Fig. 3c. Figure S6. Copy number variation of stress-related genes.

Figure S7. Patterns of differentially expressed genes between *Vigna* species (leaf).

Figure S8. Patterns of differentially expressed genes between *Vigna* species (root).

Figure S9. Linkage map of *V. trilobata*.

Table S1. Stats of sequenced reads.

Table S2. Number of Orthogroups that are common in all species, specific to Fabids, to Vigna, or to each species.

Table S3. Number of genes in Orthogroups that are common in all species, specific to Fabids, Vigna or each species, or not assigned to any Orthogroups.

Table S4. Summary of TE annotations in each assembly.

Table S5. Presence variations across Asian Vigna species.

Table S6. Enriched GO terms in species-specific genes (pan-genome).

Table S7. *sWOX* genes located within PVs in *V. riukiuensis*.

Table S8. Complete list of positively-selected genes.

Table S9. Selected salinity- and drought-related genes upregulated in tolerant species.

