## Supplementary figures for "Genome sequence of 12 *Vigna* species as a knowledge base of stress tolerance and resistance"

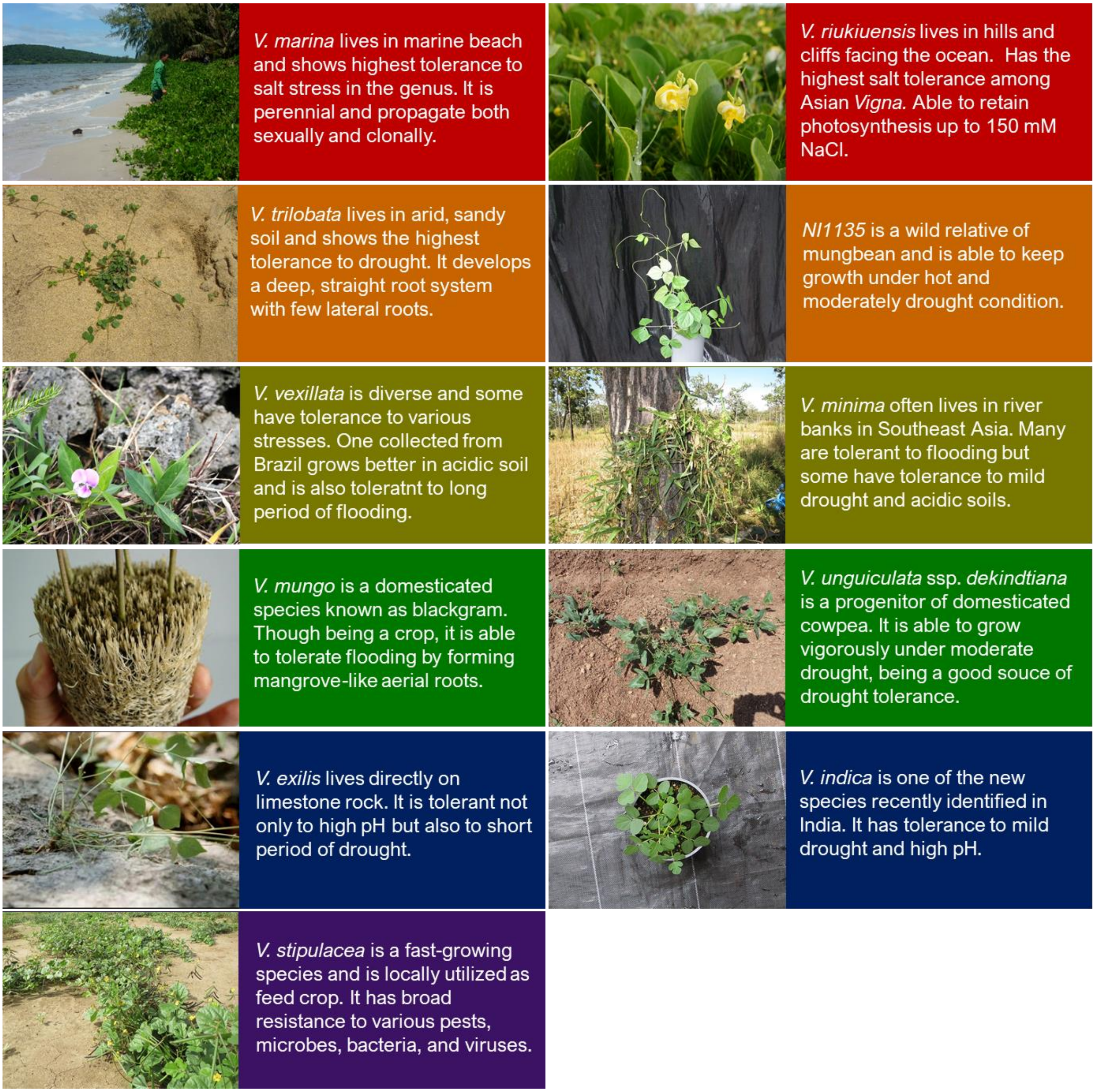


**Figure S1. Introduction of the wild species sequenced in this study.**

Fig. S2

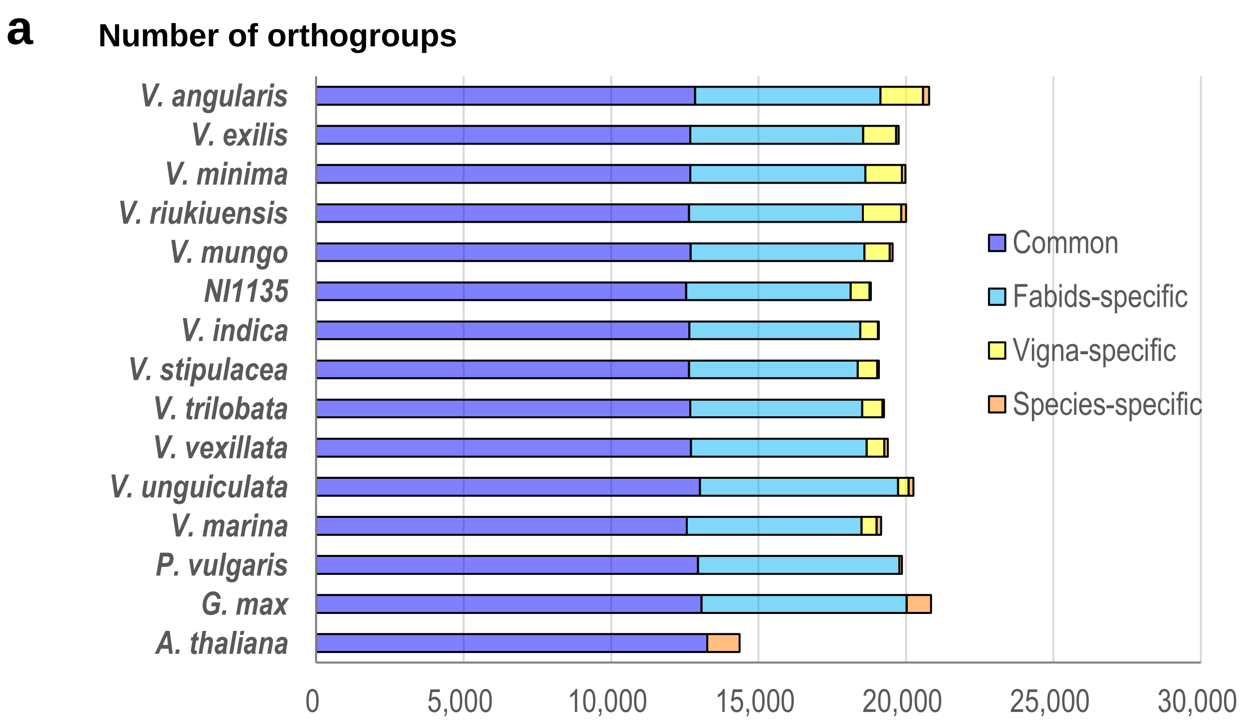


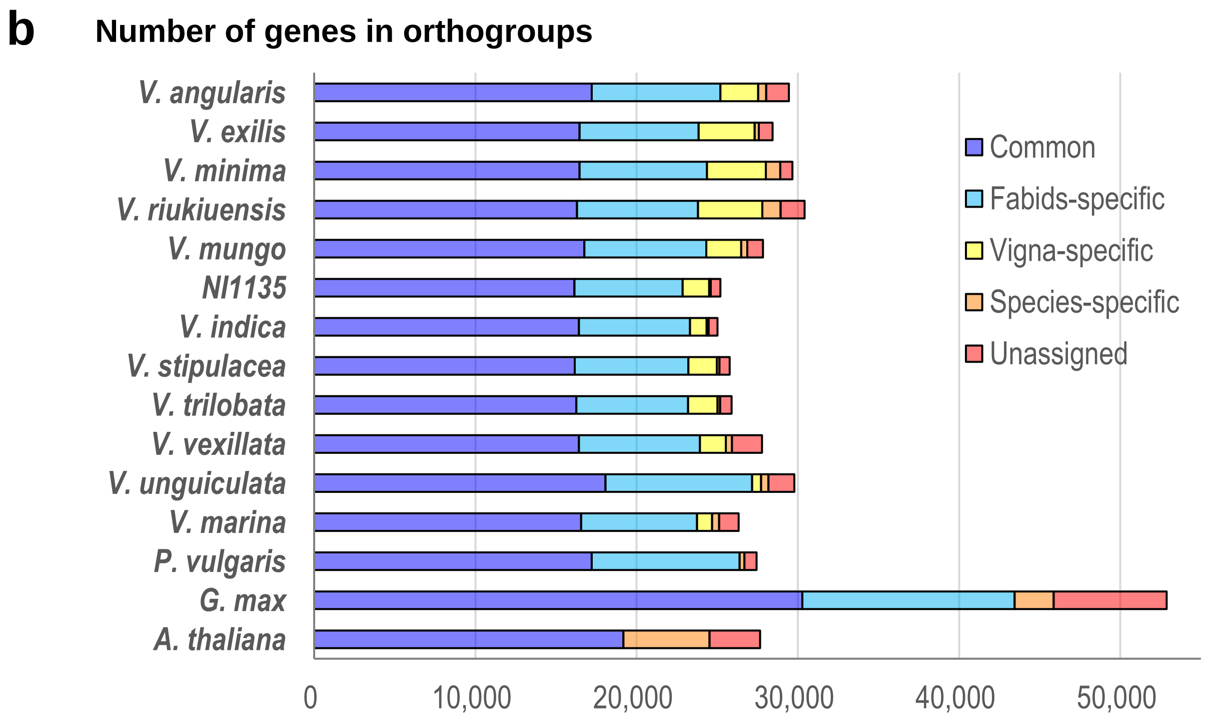


**Figure S2. Orthologous gene families in *Vigna, Phaseolus, Glycine* and *Arabidopsis.* a.** Number of orthogroups that are common in all species, specific to Fabids, *Vigna*, or each species. **b.** Number of genes in orthogroups. Colors indicate the same categories as in a. except number of genes that are not assigned to any orthogroups.

Fig. S3

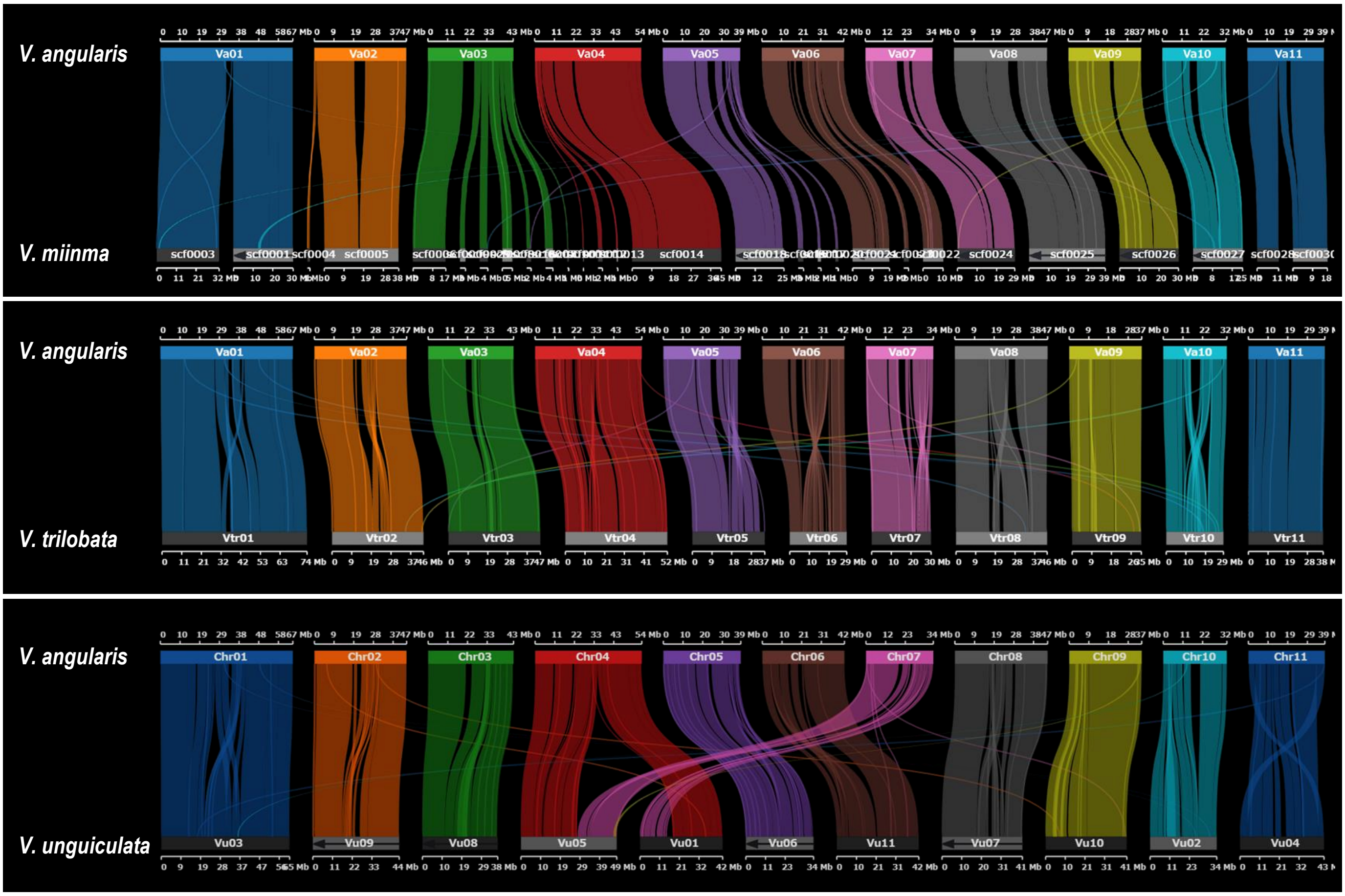


**Figure S3. Synteny plot between *V. angularis* and *V. minima*, *V. trilobata* or *V. unguiculata*.** The genome sequence of *V. unguiculata* is the one sequence d by Lonardi et al. (2019).

### Fig. S4

**a**

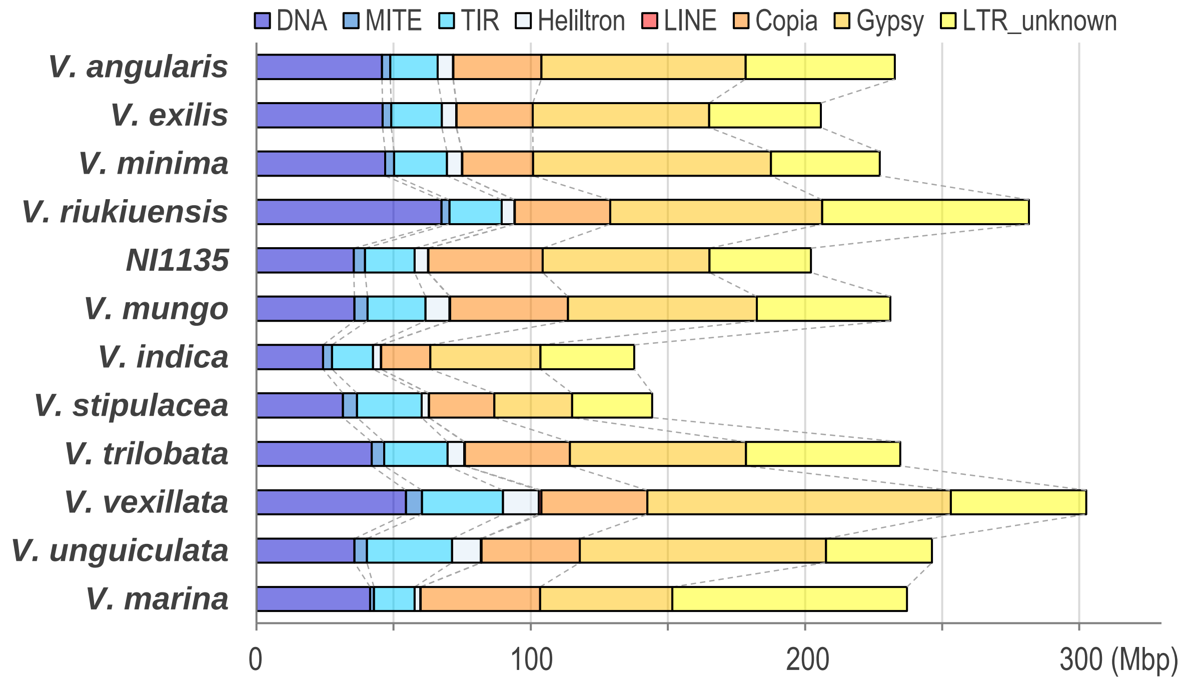


**b**

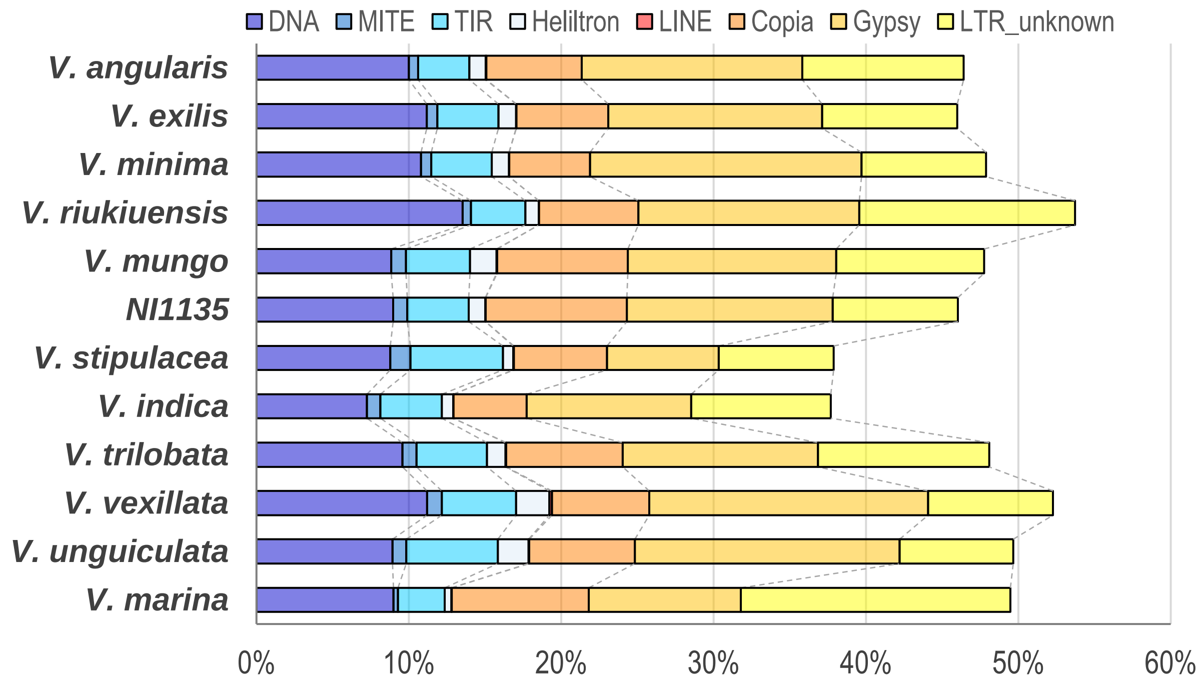


**Figure S4. TE contents in the genomes of 12 *Vigna* species. a.** Total bases comprised of TE. **b.** Percent of genomes shared by TEs.

### Fig. S5

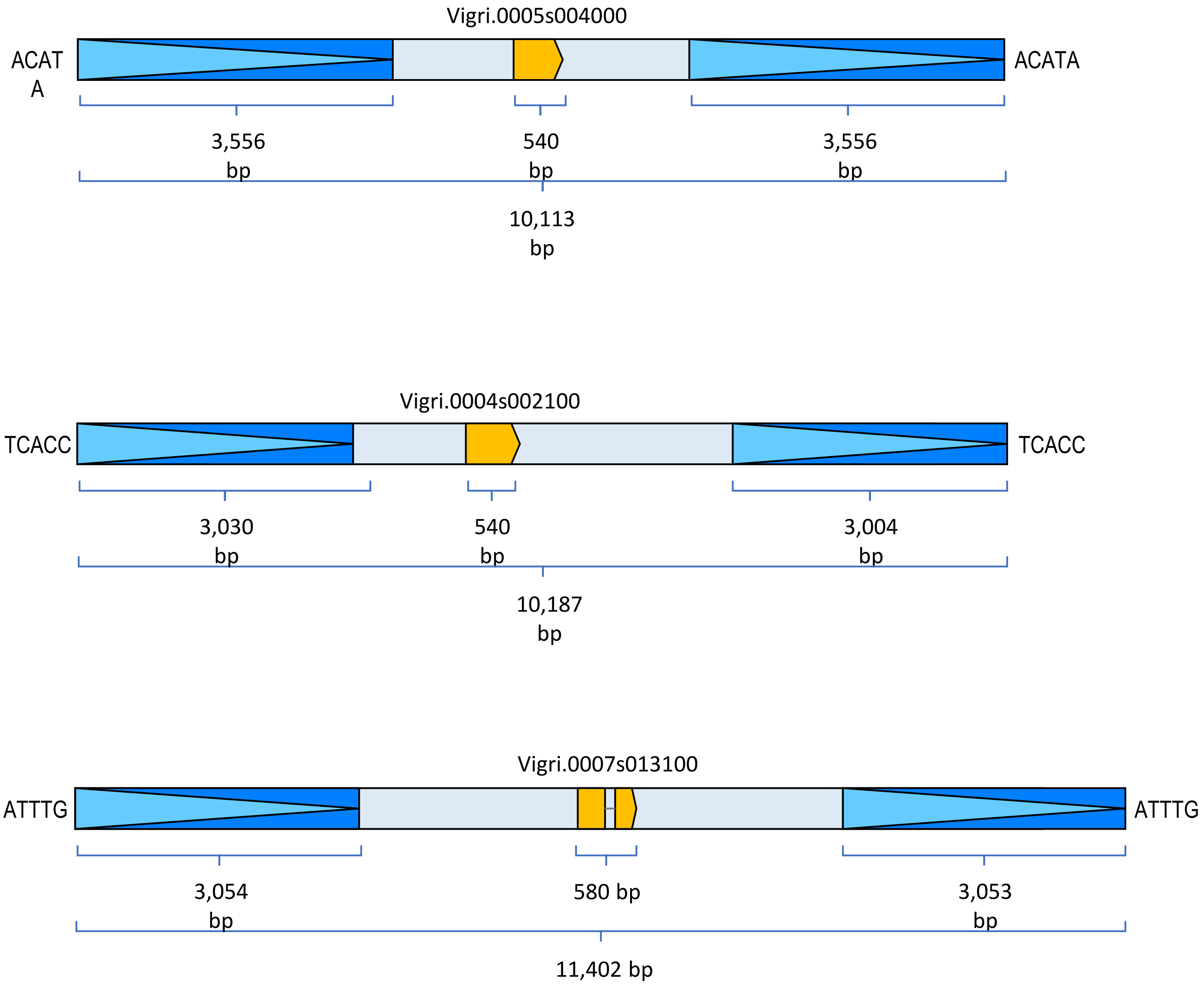


**Figure S5. Close-ups of the LTR retrotransposons and the horbored sWOX genes in Fig. 3c.**

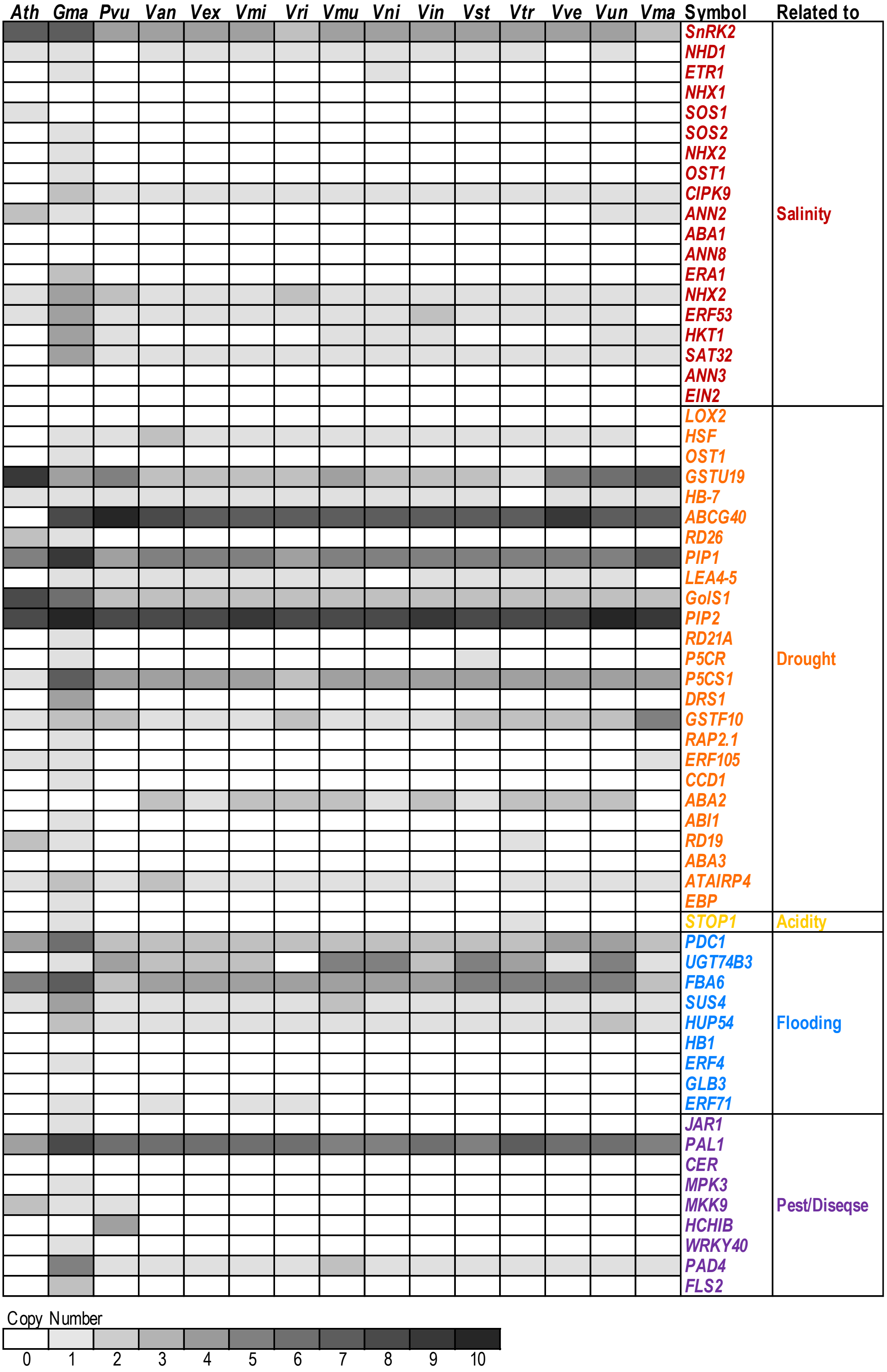


**Figure S6. Copy number variation of stress-related genes.** Well-characterized genes for tolerance to salinity (red), drought (orange), low pH (yellow), flooding/hypoxia (blue) and biotic stresses (purple) are arbitrarily selected. Graphical legend indicates copy numbers. *Ath, Gma, Pvu, Van, Vex, Vmi, Vri, Vmu, Vni, Vin, Vst, Vtr, Vve, Vun,* and *Vma* indicate *A. thaliana. G. max, P. vulgaris, V. angularis, V. exilis, V. minima, V. riukiuensis, V. mungo, NI1135, V. indica, V. stipulacea, V. trilobata, V. vexillata, V. unguiculata,* and *V. marina,* respectively.

Fig. S7

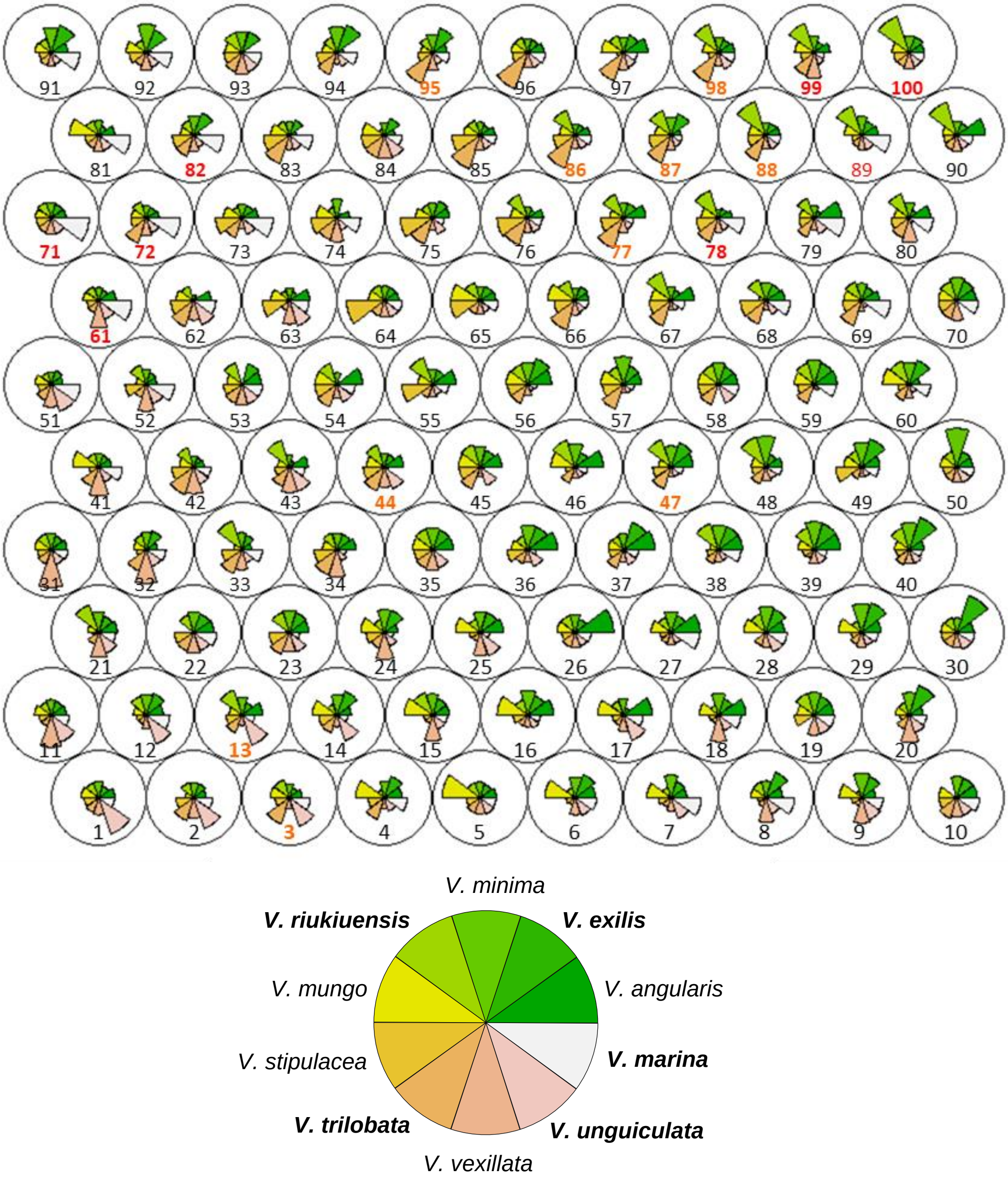


**Figure S7. Patterns of differentially expressed genes between *Vigna* species (leaf)**. The patterns are identified by SOM-clustering. Cluster numbers in red and orange indicate those with higher expression in more than 2 species of salt tolerance and drought tolerance, respectively. The height of each rose diagram indicates mean FPKM in each cluster. Species with salt tolerance (*V. marina* and *V.* riukiuensis) or drought tolerance (*V. exilis, V. riukiuensis, V. trilobata* and *V. unguiculata*) are bolded in the graphical legend.

### Fig. S8

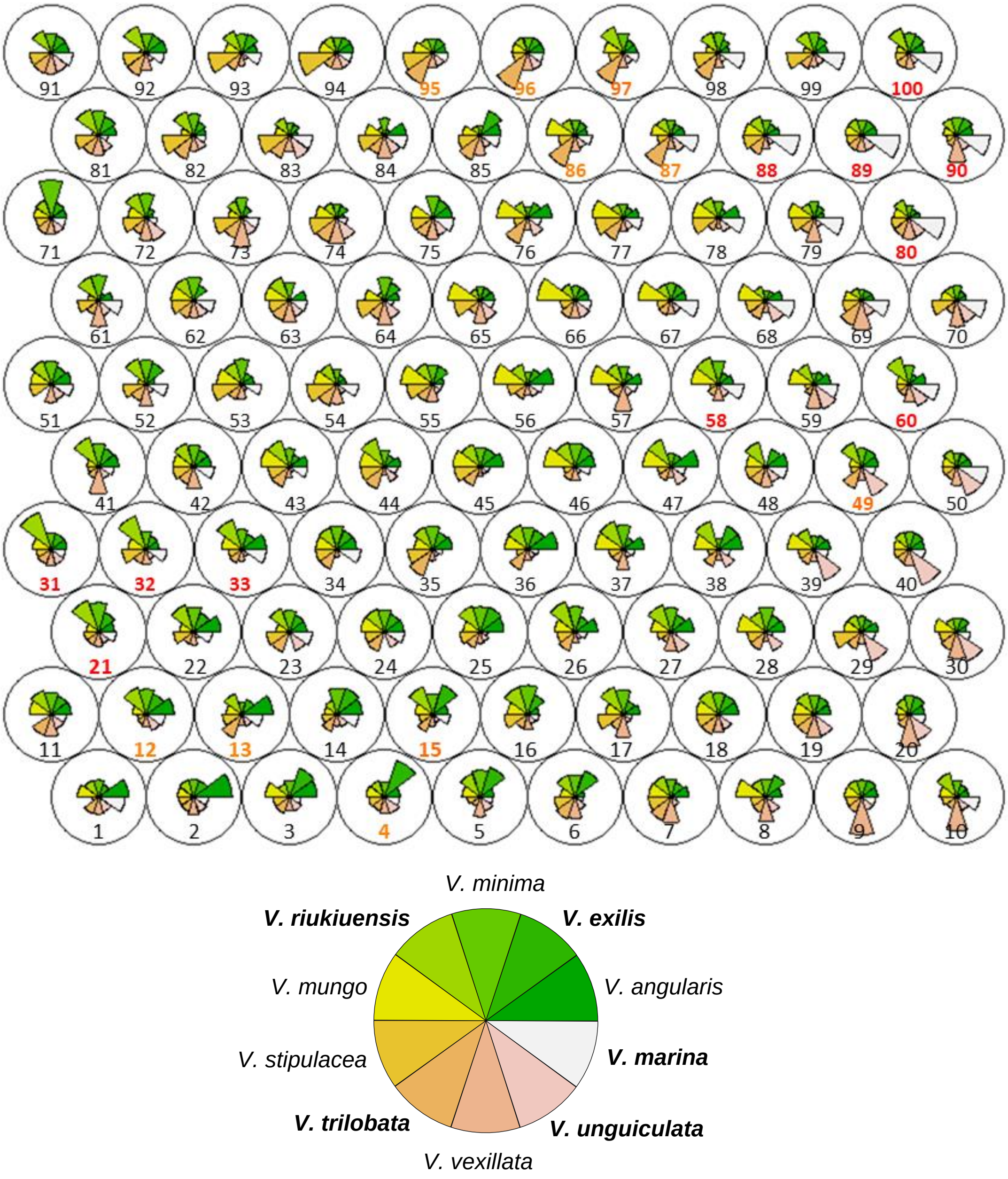


**Figure S8. Patterns of differentially expressed genes between *Vigna* species (root).** The patterns are identified by SOM-clustering. Cluster numbers in red and orange indicate those with higher expression in more than 2 species of salt tolerance and drought tolerance, respectively. The height of each rose diagram indicates mean FPKM in each cluster. Species with salt tolerance (*V. marina* and *V.* riukiuensis) or drought tolerance (*V. exilis, V. riukiuensis, V. trilobata* and *V. unguiculata*) are bolded in the graphical legend.

### Fig. S9

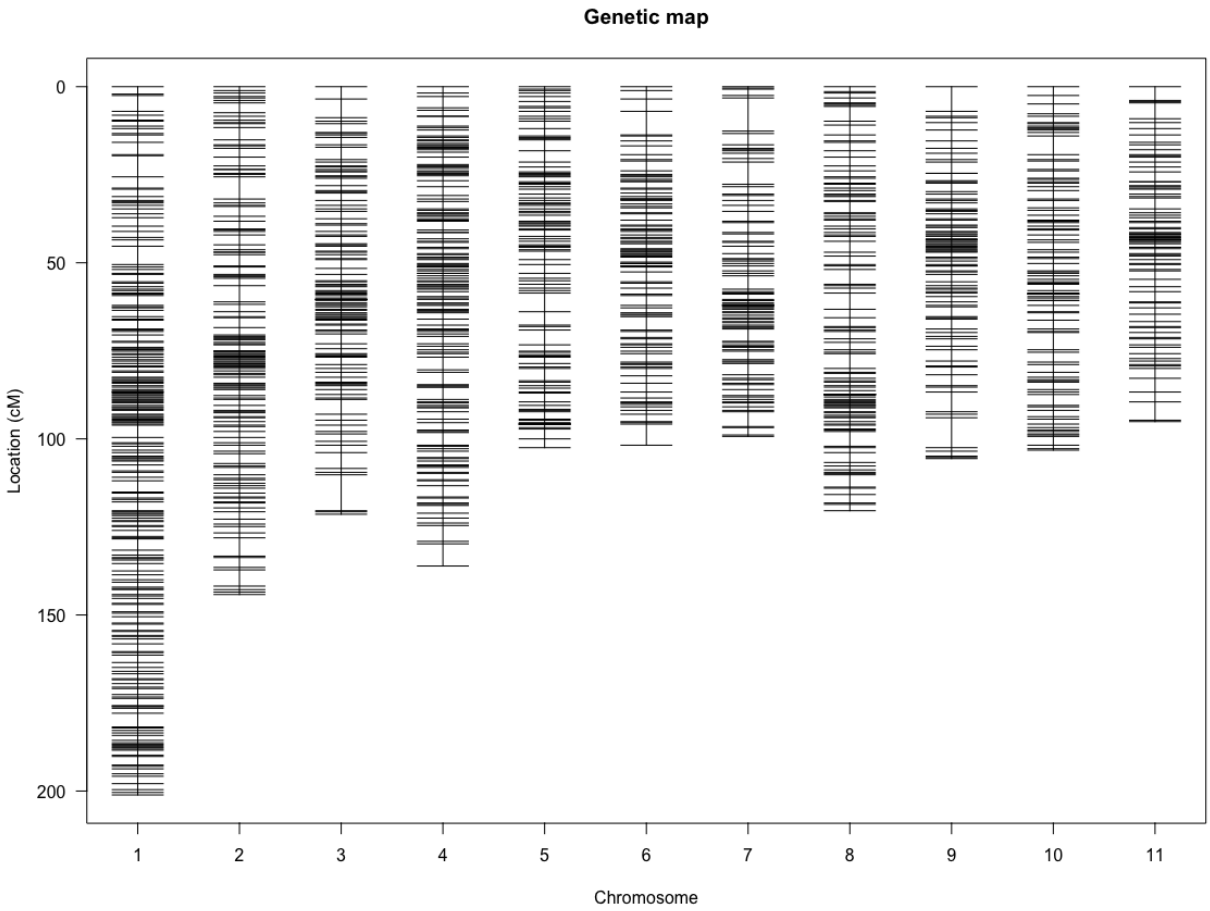


**Figure S9. Linkage map of *V. trilobata*.** We did RAD-seq analysis on the 192 F2 plants derived from JP210605 X JP252972 from RAD-seq data, genotyped and calculated genetic distance between each polymorphic loci.
