## Supplementary tables for "Genome sequence of 12 *Vigna* species as a knowledge base of stress tolerance and resistance"

Table S1. Stats of sequenced reads.

| Species | PacBio | | | Illumina_PE | | Illumina_MP3k | | Illumina_MP8k | | Illumina_MP20k | | Reference |
|---|---|---|---|---|---|---|---|---|---|---|---|---|
| | Num_reads (M) | Sum_length (Gbp) | N50_length (kbp) | Num_reads (M) | Sum_length (Gbp) | Num_reads (M) | Sum_length (Gbp) | Num_reads (M) | Sum_length (Gbp) | Num_reads (M) | Sum_length (Gbp) | |
| *V. angularis* | 6.4 | 31.3 | 7.8 | 301.6 | 45.5 | 179.7 | 16.4 | 215.2 | 19.6 | 131.1 | 12.2 | Sakai et al. 2015 |
| *V. exilis* | 2.2 | 16.4 | 15.6 | 150.6 | 22.6 | | | | | | | This study |
| *V. minima* | 1.8 | 15.5 | 14.8 | 113.5 | 11.4 | | | | | | | This study |
| *V. riukiuensis* | 5.1 | 40.6 | 11.1 | 294.7 | 44.5 | 241.1 | 21.2 | 109.7 | 9.6 | 46.6 | 4.0 | This study |
| *V. mungo* | 2.3 | 16.5 | 11.1 | 113.7 | 11.4 | | | | | | | This study |
| *NI1135* | 1.8 | 15.4 | 14.7 | 127.3 | 12.7 | | | | | | | This study |
| *V. stipulacea* | 3.5 | 20.6 | 9.5 | 102.0 | 10.3 | | | | | | | Takahashi et al. 2020 |
| *V. indica* | 4.0 | 25.8 | 11.1 | 120.1 | 12.0 | | | | | | | This study |
| *V. trilobata* | 2.5 | 16.6 | 10.0 | 294.8 | 44.5 | 397.0 | 40.1 | 289.4 | 29.2 | | | This study |
| *V. vexillata* | 2.7 | 16.3 | 9.2 | 126.9 | 12.7 | | | | | | | This study |
| *V. unguiculata* ssp. *dekindtiana* | 4.4 | 26.3 | 10.6 | | | | | | | | | Takahashi et al. 2019 |
| *V. marina* | 6.6 | 53.1 | 9.8 | 297.0 | 44.8 | 191.4 | 17.0 | 313.7 | 27.0 | 200.8 | 18.0 | This study |

Table S2. Number of Orthogroups that are common in all species, specific to Fabids, to Vigna, or to each species.

| | *V. angularis* | *V. exilis* | *V. minima* | *V. riukiuensis* | *V. mungo* | *NI1135* | *V. indica* | *V. stipulacea* | *V. trilobata* | *V. vexillata* | *V. unguiculata* | *V. marina* | *P. vulgaris* | *G. max* | *A. thaliana* |
|---|---|---|---|---|---|---|---|---|---|---|---|---|---|---|---|
| Common | 12,845 | 12,683 | 12,684 | 12,647 | 12,693 | 12,540 | 12,658 | 12,640 | 12,685 | 12,712 | 13,010 | 12,561 | 12,943 | 13,064 | 13,258 |
| Fabids-specific | 6,292 | 5,866 | 5,935 | 5,888 | 5,894 | 5,583 | 5,793 | 5,723 | 5,826 | 5,956 | 6,717 | 5,928 | 6,832 | 6,958 | 0 |
| *Vigna*-specific | 1,440 | 1,119 | 1,238 | 1,304 | 863 | 642 | 601 | 659 | 684 | 595 | 364 | 515 | 0 | 0 | 0 |
| Species-specific | 204 | 84 | 127 | 162 | 92 | 44 | 31 | 60 | 63 | 117 | 160 | 150 | 84 | 830 | 1,104 |
| Total | 20,781 | 19,752 | 19,984 | 20,001 | 19,542 | 18,809 | 19,083 | 19,082 | 19,258 | 19,380 | 20,251 | 19,154 | 19,859 | 20,852 | 14,362 |

Table S3. Number of genes in Orthogroups that are common in all species, specific to Fabids, Vigna or each species, or not assigned to any Orthogroups.

| | *V. angularis* | *V. exilis* | *V. minima* | *V. riukiuensis* | *V. mungo* | *NI1135* | *V. indica* | *V. stipulacea* | *V. trilobata* | *V. vexillata* | *V. unguiculata* | *V. marina* | *P. vulgaris* | *G. max* | *A. thaliana* |
|---|---|---|---|---|---|---|---|---|---|---|---|---|---|---|---|
| Common | 17,219 | 16,464 | 16,459 | 16,299 | 16,758 | 16,138 | 16,410 | 16,156 | 16,260 | 16,426 | 18,060 | 16,565 | 17,221 | 30,286 | 19,170 |
| Fabids-specific | 7,984 | 7,390 | 7,901 | 7,505 | 7,557 | 6,711 | 6,904 | 7,054 | 6,929 | 7,493 | 9,097 | 7,186 | 9,170 | 13,160 | 0 |
| *Vigna*-specific | 2,338 | 3,469 | 3,658 | 3,987 | 2,164 | 1,645 | 1,031 | 1,775 | 1,832 | 1,609 | 563 | 919 | 0 | 0 | 0 |
| Species-specific | 495 | 263 | 892 | 1,145 | 382 | 97 | 115 | 143 | 145 | 392 | 450 | 450 | 288 | 2,429 | 5,356 |
| Unassigned | 1,409 | 845 | 755 | 1,491 | 970 | 604 | 553 | 641 | 716 | 1,866 | 1,603 | 1,199 | 754 | 6,997 | 3,128 |
| Total | 29,445 | 28,431 | 29,665 | 30,427 | 27,831 | 25,195 | 25,013 | 25,769 | 25,882 | 27,786 | 29,773 | 26,319 | 27,433 | 52,872 | 27,654 |

**Table S4**. Summary of TE annotations in each assembly.

| **TE superfamily** | | ***V. angularis*** | | | ***V. exilis*** | | | ***V. minima*** | | | ***V. riukiuensis*** | | | ***V. mungo*** | | | ***NI1135*** | | |
|---|---|---|---|---|---|---|---|---|---|---|---|---|---|---|---|---|---|---|---|
| DNA | *hAT* | 8,539 | 3,114,295 | 0.61% | 6,646 | 2,458,031 | 0.54% | 8,784 | 2,630,518 | 0.54% | 19,393 | 4,427,495 | 0.83% | 14,906 | 4,493,364 | 0.89% | 15,266 | 4,266,465 | 0.95% |
| | *CACTA* | 45,219 | 16,353,261 | 3.19% | 44,913 | 19,236,321 | 4.19% | 44,655 | 19,190,960 | 3.95% | 55,215 | 19,255,387 | 3.61% | 34,274 | 11,914,062 | 2.37% | 26,067 | 9,584,059 | 2.13% |
| | *PIF-Harbinger* | 1,216 | 386,813 | 0.08% | 1,877 | 417,567 | 0.09% | 5,885 | 1,528,214 | 0.31% | 394 | 140,859 | 0.03% | 2,183 | 557,746 | 0.11% | 1,453 | 279,837 | 0.06% |
| | *Mutator* | 68,383 | 23,665,396 | 4.61% | 66,484 | 21,569,863 | 4.70% | 54,739 | 20,205,970 | 4.16% | 100,137 | 41,703,952 | 7.83% | 54,268 | 18,028,152 | 3.59% | 58,922 | 20,228,415 | 4.50% |
| | *Tc1-mariner* | 9,099 | 2,270,639 | 0.44% | 6,887 | 2,273,979 | 0.50% | 9,296 | 3,429,788 | 0.71% | 7,532 | 1,889,017 | 0.35% | 2,712 | 713,696 | 0.14% | 5,269 | 1,109,837 | 0.25% |
| | *Helitron* | 20,903 | 5,557,446 | 1.08% | 19,560 | 5,333,708 | 1.16% | 15,816 | 5,524,904 | 1.14% | 15,659 | 4,709,952 | 0.88% | 19,128 | 8,716,639 | 1.74% | 21,460 | 4,898,732 | 1.09% |
| LINE | *unknown* | 501 | 147,202 | 0.03% | 229 | 79,472 | 0.02% | 206 | 72,232 | 0.01% | 194 | 69,727 | 0.01% | 404 | 188,023 | 0.04% | 297 | 113,888 | 0.03% |
| LTR | *Copia* | 49,065 | 32,148,942 | 6.26% | 40,848 | 27,701,929 | 6.04% | 37,915 | 25,785,590 | 5.30% | 41,796 | 34,742,896 | 6.52% | 62,871 | 43,032,465 | 8.57% | 55,478 | 41,641,867 | 9.26% |
| | *Gypsy* | 110,292 | 74,343,395 | 14.48% | 88,567 | 64,389,869 | 14.03% | 108,665 | 86,694,415 | 17.83% | 102,009 | 77,322,303 | 14.51% | 97,613 | 68,774,981 | 13.69% | 84,728 | 60,817,579 | 13.52% |
| | *unknown* | 119,553 | 54,417,173 | 10.60% | 84,366 | 40,687,158 | 8.87% | 85,529 | 39,655,745 | 8.16% | 149,120 | 75,395,751 | 14.15% | 123,492 | 48,745,184 | 9.70% | 88,266 | 36,925,205 | 8.21% |
| MITE | *hAT* | 3,548 | 1,062,665 | 0.21% | 4,053 | 941,184 | 0.21% | 3,762 | 1,065,487 | 0.22% | 3,531 | 868,392 | 0.16% | 3,001 | 862,897 | 0.17% | 3,918 | 790,357 | 0.18% |
| | *CACTA* | 563 | 130,693 | 0.03% | 492 | 68,461 | 0.01% | 572 | 88,366 | 0.02% | 224 | 31,419 | 0.01% | 954 | 206,105 | 0.04% | 387 | 70,317 | 0.02% |
| | *PIF-Harbinger* | 1,434 | 264,237 | 0.05% | 1,257 | 296,140 | 0.06% | 1,062 | 241,247 | 0.05% | 1,542 | 316,845 | 0.06% | 846 | 185,733 | 0.04% | 822 | 167,309 | 0.04% |
| | *Mutator* | 9,441 | 1,501,229 | 0.29% | 8,279 | 1,761,002 | 0.38% | 9,374 | 1,709,463 | 0.35% | 8,689 | 1,656,908 | 0.31% | 17,776 | 3,492,471 | 0.70% | 12,757 | 3,004,268 | 0.67% |
| | *Tc1-mariner* | 709 | 108,646 | 0.02% | 756 | 124,916 | 0.03% | 786 | 104,575 | 0.02% | 770 | 132,529 | 0.02% | 661 | 84,506 | 0.02% | 364 | 52,246 | 0.01% |
| TIR | *EnSpm_CACTA* | 107 | 29,448 | 0.01% | 1,063 | 394,073 | 0.09% | 2,810 | 623,022 | 0.13% | 1,326 | 454,241 | 0.09% | 591 | 182,167 | 0.04% | 38 | 21,348 | 0.00% |
| | *MuDR_Mutator* | 956 | 315,207 | 0.06% | 255 | 112,710 | 0.02% | 240 | 183,357 | 0.04% | 221 | 119,976 | 0.02% | 450 | 244,634 | 0.05% | 577 | 411,802 | 0.09% |
| | *PIF-Harbinger* | 0 | 0 | 0 | 454 | 139,445 | 0.03% | 0 | 0 | 0 | 0 | 0 | 0 | 0 | 0 | 0 | 365 | 236,607 | 0.05% |
| | *hAT* | 0 | 0 | 0 | 72 | 32,725 | 0.01% | 0 | 0 | 0 | 0 | 0 | 0 | 146 | 48,215 | 0.01% | 0 | 0 | 0 |
| | *Unknown* | 70,437 | 16,887,255 | 3.29% | 66,049 | 17,735,590 | 3.86% | 71,356 | 18,474,242 | 3.80% | 73,485 | 18,409,079 | 3.45% | 79,585 | 20,650,962 | 4.11% | 69,877 | 17,502,555 | 3.89% |
| Total | | 519,965 | 232,703,942 | 45.33% | 443,107 | 205,754,143 | 44.84% | 461,630 | 227,275,999 | 46.74% | 581,294 | 281,671,691 | 52.86% | 515,985 | 231,203,928 | 46.03% | 446,378 | 202,189,661 | 44.95% |

**Table S4.** Summary of TE annotations in each assembly, continued.

| **TE superfamily** | | ***V. stipulacea*** | | | ***V. indica*** | | | ***V. trilobata*** | | | ***V. vexillata*** | | | ***V. unguiculata*** | | | ***V. marina*** | | |
|---|---|---|---|---|---|---|---|---|---|---|---|---|---|---|---|---|---|---|---|
| DNA | *hAT* | 7,311 | 4,337,224 | 1.12% | 8,713 | 2,342,655 | 0.63% | 10,261 | 2,863,380 | 0.57% | 24,368 | 8,079,810 | 1.34% | 23,873 | 7,436,014 | 1.44% | 12,365 | 4,748,832 | 0.98% |
| | *CACTA* | 23,845 | 7,924,150 | 2.04% | 25,743 | 8,580,724 | 2.30% | 38,410 | 12,337,682 | 2.46% | 57,692 | 20,247,478 | 3.35% | 37,694 | 13,217,289 | 2.56% | 26,602 | 10,611,761 | 2.20% |
| | *PIF-Harbinger* | 2,560 | 976,258 | 0.25% | 818 | 186,881 | 0.05% | 3,465 | 905,196 | 0.18% | 5,296 | 1,450,298 | 0.24% | 2,163 | 702,934 | 0.14% | 1,918 | 474,202 | 0.10% |
| | *Mutator* | 44,104 | 15,476,210 | 3.99% | 41,889 | 12,504,134 | 3.35% | 58,901 | 21,643,109 | 4.32% | 48,673 | 24,102,081 | 3.99% | 45,998 | 13,744,935 | 2.66% | 81,817 | 24,530,675 | 5.07% |
| | *Tc1-mariner* | 6,965 | 2,798,097 | 0.72% | 3,555 | 636,340 | 0.17% | 12,839 | 4,236,967 | 0.85% | 3,225 | 686,848 | 0.11% | 3,033 | 641,127 | 0.12% | 3,103 | 1,028,848 | 0.21% |
| | *Helitron* | 10,246 | 2,580,063 | 0.67% | 11,701 | 2,841,105 | 0.76% | 19,809 | 6,076,946 | 1.21% | 49,233 | 13,075,530 | 2.17% | 37,417 | 10,463,433 | 2.02% | 8,091 | 2,135,548 | 0.44% |
| LINE | *unknown* | 383 | 111,808 | 0.03% | 69 | 24,006 | 0.01% | 469 | 207,429 | 0.04% | 1,126 | 1,010,963 | 0.17% | 628 | 275,361 | 0.05% | 236 | 150,252 | 0.03% |
| LTR | *Copia* | 32,219 | 23,752,747 | 6.13% | 28,274 | 17,943,184 | 4.81% | 48,104 | 38,280,847 | 7.65% | 67,815 | 38,610,207 | 6.39% | 61,303 | 35,785,154 | 6.93% | 55,598 | 43,479,586 | 8.99% |
| | *Gypsy* | 36,023 | 28,404,400 | 7.33% | 76,787 | 40,239,400 | 10.79% | 88,057 | 64,180,028 | 12.82% | 181,079 | 110,603,790 | 18.31% | 139,551 | 89,764,492 | 17.37% | 72,531 | 48,179,712 | 9.97% |
| | *unknown* | 61,085 | 29,181,669 | 7.53% | 134,662 | 34,174,834 | 9.17% | 126,432 | 56,246,409 | 11.24% | 106,657 | 49,402,783 | 8.18% | 88,032 | 38,617,615 | 7.47% | 152,555 | 85,527,266 | 17.69% |
| MITE | *hAT* | 3,228 | 830,849 | 0.21% | 3,401 | 590,212 | 0.16% | 3,973 | 976,310 | 0.20% | 4,735 | 1,361,709 | 0.23% | 6,208 | 1,725,499 | 0.33% | 1,187 | 291,024 | 0.06% |
| | *CACTA* | 258 | 31,919 | 0.01% | 154 | 32,097 | 0.01% | 149 | 38,642 | 0.01% | 2,202 | 331,647 | 0.05% | 326 | 81,306 | 0.02% | 718 | 98,573 | 0.02% |
| | *PIF-Harbinger* | 530 | 94,935 | 0.02% | 15 | 3,171 | 0.00% | 506 | 107,709 | 0.02% | 186 | 39,789 | 0.01% | 4 | 1,021 | 0.00% | 62 | 11,994 | 0.00% |
| | *Mutator* | 16,635 | 4,139,460 | 1.07% | 14,072 | 2,588,180 | 0.69% | 12,039 | 3,293,914 | 0.66% | 20,805 | 4,052,912 | 0.67% | 11,796 | 2,772,640 | 0.54% | 4,642 | 1,002,987 | 0.21% |
| | *Tc1-mariner* | 152 | 28,613 | 0.01% | 238 | 41,483 | 0.01% | 1,039 | 169,056 | 0.03% | 22 | 2,730 | 0.00% | 12 | 1,155 | 0.00% | 3 | 835 | 0.00% |
| TIR | *EnSpm_CACTA* | 367 | 172,021 | 0.04% | 94 | 48,788 | 0.01% | 0 | 0 | 0 | 939 | 161,631 | 0.03% | 889 | 189,410 | 0.04% | 0 | 0 | 0 |
| | *MuDR_Mutator* | 711 | 623,951 | 0.16% | 249 | 259,704 | 0.07% | 191 | 130,966 | 0.03% | 184 | 45,250 | 0.01% | 115 | 46,128 | 0.01% | 425 | 184,547 | 0.04% |
| | *PIF-Harbinger* | 0 | 0 | 0 | 170 | 108,551 | 0.03% | 0 | 0 | 0 | 0 | 0 | 0 | 312 | 187,696 | 0.04% | 113 | 48,598 | 0.01% |
| | *hAT* | 233 | 117,368 | 0.03% | 222 | 101,847 | 0.03% | 205 | 92,299 | 0.02% | 784 | 272,649 | 0.05% | 317 | 114,250 | 0.02% | 361 | 118,258 | 0.02% |
| | *Unknown* | 83,315 | 22,692,980 | 5.85% | 55,480 | 14,511,978 | 3.89% | 86,160 | 22,939,958 | 4.58% | 113,981 | 29,007,162 | 4.80% | 108,837 | 30,468,738 | 5.90% | 53,768 | 14,499,367 | 3.00% |
| Total | | 330,227 | 144,321,089 | 37.22% | 406,407 | 137,822,904 | 36.97% | 511,340 | 235,086,665 | 46.96% | 689,945 | 302,857,810 | 50.15% | 569,398 | 246,834,092 | 47.77% | 476,630 | 237,438,629 | 49.12% |

**Table S5. Presence variations across Asian Vigna species.**

| species | Number of PVs | Toltal length (Mbp) | Mean length (bp) | Median length (bp) | Max. length (bp) |
|---|---|---|---|---|---|
| *V. angularis* | 3,299 | 12.47 | 3,778.6 | 2,541.0 | 72,082.0 |
| *V. exilis* | 2,590 | 8.25 | 3,184.8 | 2,265.5 | 41,373.0 |
| *V. minima* | 2,564 | 9.32 | 3,634.4 | 2,455.5 | 47,676.0 |
| *V. riukiuensis* | 4,206 | 20.64 | 4,906.5 | 3,454.0 | 59,787.0 |
| *V. mungo* | 3,670 | 11.66 | 3,175.9 | 2,276.0 | 44,965.0 |
| *NI1135* | 3,074 | 8.64 | 2,811.4 | 1,620.0 | 38,425.0 |
| *V. indica* | 2,061 | 4.95 | 2,403.4 | 1,383.0 | 34,564.0 |
| *V. stipulacea* | 2,379 | 6.13 | 2,578.3 | 1,411.0 | 28,508.0 |
| *V. trilobata* | 3,504 | 11.76 | 3,356.2 | 2,198.0 | 51,701.0 |

**Table S6. Enriched GO terms in species-specific genes (pan-genome).**

| | | | | Specific genes | | | Other genes | | | | |
|---|---|---|---|---|---|---|---|---|---|---|---|
| ***Species*** | **GOID** | **Categ** | **Description** | **Hit** | **No-** | **Hit/No-** | **Hit** | **No-** | **Hit/No-** | **Rati** | **pvalue** |
| *V. angularis* | 51315 | BP | attachment of mitotic spindle | 4 | 1900 | 0.002 | 1 | 27540 | 0 | 58 | 0.019 |
| ***V. marina*** | **3700** | **MF** | **DNA-binding transcription** | **88** | **1561** | **0.056** | **648** | **24119** | **0.026** | **2.1** | **9.48E-07** |
| ***V. marina*** | **99402** | **BP** | **plant organ development** | **58** | **1591** | **0.036** | **13** | **24754** | **0.001** | **69.4** | **3.02E-55** |
| *V. minima* | 47834 | MF | D-threo-aldose 1- | 15 | 1632 | 0.009 | 41 | 28106 | 0.001 | 6.3 | 2.98E-05 |
| ***V. riukiuensis*** | **3677** | **MF** | **DNA binding** | **829** | **1807** | **0.458** | **1112** | **26750** | **0.041** | **11** | **0** |
| ***V. riukiuensis*** | **3700** | **MF** | **DNA-binding transcription** | **807** | **1829** | **0.441** | **811** | **27051** | **0.029** | **14.7** | **0** |
| ***V. riukiuensis*** | **99402** | **BP** | **plant organ development** | **583** | **2053** | **0.283** | **95** | **27767** | **0.003** | **83** | **0** |
| *V. stipulacea* | 4489 | MF | methylenetetrahydrofolate | 3 | 781 | 0.003 | 1 | 25253 | 0 | 97 | 0.014 |
| *V. stipulacea* | 6555 | BP | methionine metabolic process | 3 | 781 | 0.003 | 1 | 25253 | 0 | 97 | 0.014 |
| *V. stipulacea* | 10265 | BP | SCF complex assembly | 4 | 780 | 0.005 | 4 | 25250 | 0 | 32.4 | 0.007 |
| *V. unguiculata* | 4535 | MF | poly(A)-specific ribonuclease | 9 | 749 | 0.012 | 6 | 31184 | 0 | 62.5 | 1.54E-09 |
| *V. unguiculata* | 5199 | MF | structural constituent of cell | 4 | 754 | 0.005 | 4 | 31186 | 0 | 41.4 | 0.003 |
| *V. unguiculata* | 9512 | CC | cytochrome b6f complex | 5 | 753 | 0.006 | 19 | 31171 | 0 | 10.9 | 0.033 |
| *V. unguiculata* | 30014 | CC | CCR4-NOT complex | 9 | 749 | 0.012 | 12 | 31178 | 0 | 31.2 | 7.97E-08 |
| *V. unguiculata* | 31361 | CC | integral component of thylakoid | 5 | 753 | 0.006 | 10 | 31180 | 0 | 20.7 | 0.003 |

**Table S7. sWOX genes located within PVs in *V. riukiuensis*.**

| Scaffold | PV_start | PV_end | PV_length | Gene_id |
|---|---|---|---|---|
| scf0005 | 23,023 | 32,760 | 9,738 | Vigri.0005s000100.01 |
| scf0005 | 408,475 | 418,642 | 10,168 | Vigri.0005s004000.01 |
| scf0013 | 3,638,420 | 3,664,518 | 26,099 | Vigri.0013s015900.01 |
| scf0020 | 503,497 | 510,720 | 7,224 | Vigri.0020s004300.01 |
| scf0024 | 3,548,785 | 3,558,262 | 9,478 | Vigri.0024s019500.01 |
| scf0025 | 181,216 | 191,083 | 9,868 | Vigri.0025s000900.01 |
| scf0027 | 1,556,366 | 1,565,603 | 9,238 | Vigri.0027s009300.01 |
| scf0027 | 1,696,877 | 1,706,077 | 9,201 | Vigri.0027s009800.01 |
| scf0027 | 1,823,521 | 1,851,346 | 27,826 | Vigri.0027s011100.01 |
| scf0029 | 255,651 | 272,798 | 17,148 | Vigri.0029s002100.01 |
| scf0029 | 348,076 | 356,958 | 8,883 | Vigri.0029s003000.01 |
| scf0033 | 309,003 | 318,409 | 9,407 | Vigri.0033s002200.01 |
| scf0034 | 716,613 | 728,625 | 12,013 | Vigri.0034s005200.01 |
| scf0034 | 884,144 | 895,974 | 11,831 | Vigri.0034s006800.01 |
| scf0035 | 1,330,657 | 1,340,804 | 10,148 | Vigri.0035s011600.01 |
| scf0038 | 316,366 | 326,026 | 9,661 | Vigri.0038s003600.01 |
| scf0043 | 1,947,700 | 1,960,249 | 12,550 | Vigri.0043s013200.01 |
| scf0043 | 2,716,529 | 2,731,512 | 14,984 | Vigri.0043s017800.01 |
| scf0043 | 2,766,577 | 2,771,463 | 4,887 | Vigri.0043s018200.01 |
| scf0044 | 221,110 | 230,503 | 9,394 | Vigri.0044s000700.01 |
| scf0045 | 851,868 | 853,599 | 1,732 | Vigri.0045s004600.01 |
| scf0047 | 277,387 | 288,726 | 11,340 | Vigri.0047s001900.01 |
| scf0047 | 329,231 | 337,312 | 8,082 | Vigri.0047s002100.01 |
| scf0052 | 565,808 | 577,354 | 11,547 | Vigri.0052s002600.01 |
| scf0056 | 6,127,749 | 6,135,875 | 8,127 | Vigri.0056s031200.01 |
| scf0056 | 10,135,613 | 10,146,273 | 10,661 | Vigri.0056s057400.01 |
| scf0058 | 2,749,915 | 2,757,905 | 7,991 | Vigri.0058s024700.01 |
| scf0061 | 2,194,012 | 2,216,695 | 22,684 | Vigri.0061s010100.01 |
| scf0062 | 878,514 | 893,088 | 14,575 | Vigri.0062s004900.01 |
| scf0062 | 2,589,018 | 2,599,655 | 10,638 | Vigri.0062s013800.01 |
| scf0063 | 4,680,225 | 4,704,275 | 24,051 | Vigri.0063s043100.01 |
| scf0064 | 1,197,332 | 1,216,632 | 19,301 | Vigri.0064s004300.01 |
| scf0064 | 9,474,697 | 9,485,353 | 10,657 | Vigri.0064s033500.01 |
| scf0064 | 11,490,764 | 11,498,858 | 8,095 | Vigri.0064s042900.01 |
| scf0067 | 1,850,880 | 1,879,988 | 29,109 | Vigri.0067s012500.01 |
| scf0067 | 1,850,880 | 1,879,988 | 29,109 | Vigri.0067s012700.01 |
| scf0067 | 1,982,586 | 1,991,039 | 8,454 | Vigri.0067s013500.01 |
| scf0067 | 2,480,756 | 2,490,468 | 9,713 | Vigri.0067s018500.01 |
| scf0067 | 3,347,262 | 3,367,242 | 19,981 | Vigri.0067s026400.01 |
| scf0067 | 3,347,262 | 3,367,242 | 19,981 | Vigri.0067s026500.01 |
| scf0067 | 4,720,896 | 4,737,617 | 16,722 | Vigri.0067s038400.01 |
| scf0070 | 394,779 | 399,408 | 4,630 | Vigri.0070s004400.01 |
| scf0071 | 1,133,397 | 1,154,589 | 21,193 | Vigri.0071s006900.01 |
| scf0071 | 1,133,397 | 1,154,589 | 21,193 | Vigri.0071s007000.01 |
| scf0071 | 1,133,397 | 1,154,589 | 21,193 | Vigri.0071s007100.01 |
| scf0071 | 1,971,187 | 1,980,652 | 9,466 | Vigri.0071s016100.01 |
| scf0072 | 2,895,042 | 2,904,168 | 9,127 | Vigri.0072s024500.01 |
| scf0072 | 4,883,582 | 4,893,741 | 10,160 | Vigri.0072s042900.01 |
| scf0072 | 9,201,146 | 9,218,675 | 17,530 | Vigri.0072s076600.01 |
| scf0074 | 825,573 | 834,891 | 9,319 | Vigri.0074s005800.01 |
| scf0074 | 2,004,035 | 2,014,739 | 10,705 | Vigri.0074s011900.01 |

**Table S7. sWOX genes located within PVs in *V. riukiuensis*, continued.**

| Scaffold | PV_start | PV_end | PV_length | Gene_id |
|---|---|---|---|---|
| scf0074 | 2,729,642 | 2,751,108 | 21,467 | Vigri.0074s016200.01 |
| scf0075 | 4,484,107 | 4,493,187 | 9,081 | Vigri.0075s020900.01 |
| scf0079 | 403,905 | 416,236 | 12,332 | Vigri.0079s002200.01 |
| scf0079 | 403,905 | 416,236 | 12,332 | Vigri.0079s002300.01 |
| scf0084 | 2,234,511 | 2,245,468 | 10,958 | Vigri.0084s009300.01 |
| scf0089 | 473,660 | 481,747 | 8,088 | Vigri.0089s003100.01 |
| scf0089 | 734,106 | 771,282 | 37,177 | Vigri.0089s004500.01 |
| scf0093 | 1,254,378 | 1,260,938 | 6,561 | Vigri.0093s015500.01 |
| scf0093 | 1,641,780 | 1,661,024 | 19,245 | Vigri.0093s021300.01 |
| scf0093 | 1,641,780 | 1,661,024 | 19,245 | Vigri.0093s021400.01 |
| scf0093 | 1,795,620 | 1,821,092 | 25,473 | Vigri.0093s022800.01 |
| scf0093 | 1,795,620 | 1,821,092 | 25,473 | Vigri.0093s022900.01 |
| scf0093 | 3,017,211 | 3,026,270 | 9,060 | Vigri.0093s037400.01 |
| scf0095 | 1,474,474 | 1,486,032 | 11,559 | Vigri.0095s013700.01 |
| scf0095 | 1,730,613 | 1,740,280 | 9,668 | Vigri.0095s015800.01 |
| scf0095 | 2,312,405 | 2,323,482 | 11,078 | Vigri.0095s020700.01 |
| scf0095 | 3,269,323 | 3,278,327 | 9,005 | Vigri.0095s025600.01 |
| scf0096 | 2,065,035 | 2,078,776 | 13,742 | Vigri.0096s019800.01 |
| scf0096 | 2,065,035 | 2,078,776 | 13,742 | Vigri.0096s019900.01 |
| scf0096 | 6,164,216 | 6,173,321 | 9,106 | Vigri.0096s054300.01 |
| scf0096 | 6,537,462 | 6,547,944 | 10,483 | Vigri.0096s057600.01 |
| scf0096 | 6,624,656 | 6,635,291 | 10,636 | Vigri.0096s058000.01 |
| scf0098 | 2,702,397 | 2,709,192 | 6,796 | Vigri.0098s015200.01 |
| scf0098 | 3,812,156 | 3,820,212 | 8,057 | Vigri.0098s018800.01 |
| scf0099 | 1,356,556 | 1,372,982 | 16,427 | Vigri.0099s004200.01 |
| scf0100 | 245,015 | 251,280 | 6,266 | Vigri.0100s002600.01 |
| scf0102 | 330,890 | 341,232 | 10,343 | Vigri.0102s002900.01 |
| scf0102 | 918,895 | 927,679 | 8,785 | Vigri.0102s006700.01 |
| scf0102 | 1,102,358 | 1,111,185 | 8,828 | Vigri.0102s008100.01 |
| scf0106 | 34,920 | 44,881 | 9,962 | Vigri.0106s000300.01 |
| scf0106 | 2,781,044 | 2,790,651 | 9,608 | Vigri.0106s033400.01 |
| scf0106 | 5,716,985 | 5,727,147 | 10,163 | Vigri.0106s064900.01 |
| scf0107 | 272,938 | 292,368 | 19,431 | Vigri.0107s001800.01 |
| scf0107 | 514,343 | 522,019 | 7,677 | Vigri.0107s003200.01 |
| scf0107 | 1,580,858 | 1,590,694 | 9,837 | Vigri.0107s010100.01 |
| scf0107 | 2,778,466 | 2,786,383 | 7,918 | Vigri.0107s017200.01 |
| scf0107 | 7,444,469 | 7,454,294 | 9,826 | Vigri.0107s039100.01 |
| scf0109 | 5,685,336 | 5,696,460 | 11,125 | Vigri.0109s022700.01 |
| scf0113 | 2,852,754 | 2,861,676 | 8,923 | Vigri.0113s013800.01 |
| scf0114 | 3,121,983 | 3,131,904 | 9,922 | Vigri.0114s034500.01 |
| scf0115 | 344,717 | 361,944 | 17,228 | Vigri.0115s002500.01 |
| scf0116 | 278,641 | 307,728 | 29,088 | Vigri.0116s002000.01 |
| scf0116 | 278,641 | 307,728 | 29,088 | Vigri.0116s002100.01 |
| scf0116 | 278,641 | 307,728 | 29,088 | Vigri.0116s002200.01 |
| scf0116 | 12,800,707 | 12,811,708 | 11,002 | Vigri.0116s061100.01 |
| scf0119 | 2,104,989 | 2,115,845 | 10,857 | Vigri.0119s013800.01 |
| scf0119 | 2,516,184 | 2,527,024 | 10,841 | Vigri.0119s016500.01 |
| scf0120 | 548,898 | 558,354 | 9,457 | Vigri.0120s005800.01 |
| scf0121 | 305,838 | 315,822 | 9,985 | Vigri.0121s002900.01 |
| scf0121 | 386,555 | 396,951 | 10,397 | Vigri.0121s003200.01 |

**Table S7. sWOX genes located within PVs in *V. riukiuensis*, contin**

| Scaffold | PV_start | PV_end | PV_length | Gene_id |
|---|---|---|---|---|
| scf0121 | 604,157 | 612,450 | 8,294 | Vigri.0121s004600.01 |
| scf0122 | 1,035,800 | 1,045,240 | 9,441 | Vigri.0122s010600.01 |
| scf0124 | 354,187 | 363,441 | 9,255 | Vigri.0124s003600.01 |
| scf0124 | 2,445,137 | 2,454,867 | 9,731 | Vigri.0124s016700.01 |
| scf0124 | 4,726,301 | 4,736,621 | 10,321 | Vigri.0124s035200.01 |
| scf0124 | 5,220,477 | 5,231,234 | 10,758 | Vigri.0124s039600.01 |
| scf0125 | 5,262,235 | 5,272,104 | 9,870 | Vigri.0125s040000.01 |
| scf0126 | 487,955 | 501,569 | 13,615 | Vigri.0126s003300.01 |
| scf0128 | 1,520,250 | 1,529,829 | 9,580 | Vigri.0128s006500.01 |
| scf0133 | 1,438,841 | 1,451,061 | 12,221 | Vigri.0133s009200.01 |
| scf0134 | 1,948,451 | 1,959,979 | 11,529 | Vigri.0134s006900.01 |
| scf0138 | 382,372 | 391,829 | 9,458 | Vigri.0138s002500.01 |
| scf0140 | 2,807,222 | 2,813,526 | 6,305 | Vigri.0140s031400.01 |
| scf0140 | 3,324,095 | 3,333,999 | 9,905 | Vigri.0140s035700.01 |
| scf0140 | 4,739,938 | 4,752,006 | 12,069 | Vigri.0140s046500.01 |
| scf0140 | 4,777,294 | 4,785,932 | 8,639 | Vigri.0140s046700.01 |
| scf0140 | 5,003,733 | 5,016,220 | 12,488 | Vigri.0140s047700.01 |
| scf0143 | 592,691 | 629,370 | 36,680 | Vigri.0143s002300.01 |
| scf0144 | 922,895 | 924,588 | 1,694 | Vigri.0144s004400.01 |
| scf0146 | 224,081 | 235,614 | 11,534 | Vigri.0146s000800.01 |
| scf0149 | 1,096,777 | 1,105,048 | 8,272 | Vigri.0149s004800.01 |
| scf0151 | 1,255,042 | 1,259,116 | 4,075 | Vigri.0151s007700.01 |
| scf0151 | 1,898,649 | 1,907,093 | 8,445 | Vigri.0151s011500.01 |
| scf0152 | 828,272 | 831,907 | 3,636 | Vigri.0152s005000.01 |
| scf0154 | 3,037,853 | 3,046,383 | 8,531 | Vigri.0154s024500.01 |
| scf0156 | 574,890 | 585,328 | 10,439 | Vigri.0156s003400.01 |
| scf0156 | 1,047,250 | 1,056,403 | 9,154 | Vigri.0156s004800.01 |
| scf0157 | 356,067 | 376,068 | 20,002 | Vigri.0157s001600.01 |
| scf0158 | 1,115,600 | 1,175,386 | 59,787 | Vigri.0158s005500.01 |
| scf0159 | 1,982,814 | 1,991,675 | 8,862 | Vigri.0159s014200.01 |
| scf0163 | 7,161,541 | 7,170,944 | 9,404 | Vigri.0163s055900.01 |
| scf0164 | 1,390,633 | 1,396,392 | 5,760 | Vigri.0164s005700.01 |
| scf0165 | 1,680,084 | 1,689,783 | 9,700 | Vigri.0165s007000.01 |
| scf0167 | 651,531 | 675,130 | 23,600 | Vigri.0167s006400.01 |
| scf0167 | 651,531 | 675,130 | 23,600 | Vigri.0167s006500.01 |
| scf0168 | 731,847 | 740,906 | 9,060 | Vigri.0168s003000.01 |
| scf0170 | 775,009 | 792,613 | 17,605 | Vigri.0170s004400.01 |
| scf0170 | 775,009 | 792,613 | 17,605 | Vigri.0170s004500.01 |
| scf0171 | 566,388 | 581,536 | 15,149 | Vigri.0171s002100.01 |
| scf0171 | 566,388 | 581,536 | 15,149 | Vigri.0171s002200.01 |
| scf0171 | 1,139,265 | 1,144,457 | 5,193 | Vigri.0171s005300.01 |
| scf0235 | 90,345 | 99,200 | 8,856 | Vigri.0235s000200.01 |
| scf0238 | 21,024 | 45,828 | 24,805 | Vigri.0238s000300.01 |
| scf0277 | 102,728 | 112,025 | 9,298 | Vigri.0277s001400.01 |
| scf0292 | 21,287 | 33,722 | 12,436 | Vigri.0292s000200.01 |
| scf0305 | 28,133 | 37,236 | 9,104 | Vigri.0305s000200.01 |
| scf0361 | 45,891 | 58,938 | 13,048 | Vigri.0361s000300.01 |
| scf0398 | 55,674 | 79,894 | 24,221 | Vigri.0398s000400.01 |
| scf0709 | 15,360 | 26,696 | 11,337 | Vigri.0709s000300.01 |
| scf0709 | 66,371 | 75,667 | 9,297 | Vigri.0709s000700.01 |

**Table S8. Complete list of positively-selected genes.**

| Species | Gene ID | Adjusted p-value | Closest Arabidpsis gene | Gene name | Description |
|---|---|---|---|---|---|
| *V. angularis* | Vigan.01G357900.01 | 7.17E-07 | AT4G37560.1 | IAMHYDROLASE12 (IAMH2) | Acetamidase/Formamidase family protein;(source:Araport11) |
| *V. angularis* | Vigan.01G330900.01 | 8.58E-07 | AT1G27660.1 | - | basic helix-loop-helix (bHLH) DNA-binding superfamily protein;(source:Araport11) |
| *V. angularis* | Vigan.03G018400.01 | 6.01E-05 | AT5G55710.1 | translocon at the inner envelope membrane of chloroplasts 20-V (Tic20-V) | TIC 20-v-like protein;(source:Araport11) |
| *V. exilis* | Vigex.0012s119400.01 | 0.00037 | AT4G40080.1 | PICALM10a | ENTH/ANTH/VHS superfamily protein;(source:Araport11) |
| *V. exilis* | Vigex.0007s024300.01 | 0.00329 | AT1G20760.1 | Epoxide hydrolase 1 (AtEH1) | Calcium-binding EF hand family protein;(source:Araport11) |
| *V. minima* | Vigmi.0003s002200.01 | 1.83E-06 | AT2G22475.1 | GL2-EXPRESSION MODULATOR (GEM) | GRAM domain family protein;(source:Araport11) |
| *V. mungo* | Vigmu.0717s000100.01 | 4.26E-14 | AT4G01210.1 | - | glycosyl transferase family 1 protein;(source:Araport11) |
| *V. mungo* | Vigmu.2161s000200.01 | 4.13E-08 | AT5G20850.1 | RAD51 | RAS associated with diabetes protein 51;(source:Araport11) |
| *V. mungo* | Vigmu.0081s011200.01 | 8.44E-06 | AT3G17860.1 | jasmonate-zim-domain protein 3 (JAZ3) | jasmonate-zim-domain protein 3;(source:Araport11) |
| *V. mungo* | Vigmu.0075s054800.01 | 0.00018 | AT4G01280.1 | REVEILLE 5 (RVE5) | Homeodomain-like superfamily protein;(source:Araport11) |
| *V. mungo* | Vigmu.0050s050200.01 | 0.00093 | AT5G22750.1 | RAD5 | DNA/RNA helicase protein;(source:Araport11) |
| *V. mungo* | Vigmu.0087s088100.01 | 0.01314 | AT5G45310.1 | - | coiled-coil protein;(source:Araport11) |
| *NI1135* | ni1135.0033s189000.01 | 0.00013 | AT3G09860.1 | - | actin T1-like protein;(source:Araport11) |
| *V. indica* | Vigin.0010s102300.01 | 9.44E-05 | AT5G58760.1 | damaged DNA binding 2 (DDB2) | damaged DNA binding 2;(source:Araport11) |
| *V. indica* | Vigin.0014s075000.01 | 0.00075 | AT5G66980.1 | - | AP2/B3-like transcriptional factor family protein;(source:Araport11) |
| *V. stipulacea* | Vigst.0016s062100.01 | 8.19E-09 | AT3G03440.1 | - | ARM repeat superfamily protein;(source:Araport11) |
| *V. stipulacea* | Vigst.0004s008000.01 | 1.17E-07 | AT2G37230.1 | Ribosomal Pentatricopeptide Repeat Protein 5 (RPPR5) | Tetratricopeptide repeat (TPR)-like superfamily protein;(source:Araport11) |
| *V. stipulacea* | Vigst.0008s077700.01 | 9.22E-05 | AT3G25400.1 | - | dCTP pyrophosphatase-like protein;(source:Araport11) |
| *V. trilobata* | Vigtr.08G226300.01 | 7.28E-06 | AT2G05320.1 | N-acetylglucosaminyltransferase II (GNT-II) | beta-1,2-N-acetylglucosaminyltransferase II;(source:Araport11) |
| *V. vexillata* | Vigve.0044s081100.01 | 2.38E-07 | AT2G21720.1 | - | ArgH (DUF639);(source:Araport11) |
| *V. vexillata* | Vigve.0105s061600.01 | 1.00E-06 | AT3G15790.3 | methyl-CPG-binding domain 11 (MBD11) | methyl-CPG-binding domain 11;(source:Araport11) |
| *V. unguiculata* | Vigun01g216500.1 | 1.62E-09 | AT5G37790.1 | - | Protein kinase superfamily protein;(source:Araport11) |
| *V. unguiculata* | Vigun01g000500.1 | 2.63E-08 | AT1G35510.1 | - | O-fucosyltransferase family protein;(source:Araport11) |
| *V. unguiculata* | Vigun09g150400.1 | 4.77E-08 | AT5G53450.1 | OBP3-responsive gene 1 (ORG1) | OBP3-responsive protein 1;(source:Araport11) |
| *V. unguiculata* | Vigun05g169000.1 | 0.00172 | AT1G16570.1 | TURAN (TUN) | UDP-Glycosyltransferase superfamily protein;(source:Araport11) |
| *V. unguiculata* | Vigun02g123200.1 | 0.00244 | AT5G45600.2 | GLIOMAS 41 (GAS41) | YEATS family protein;(source:Araport11) |
| *V. unguiculata* | Vigun11g117200.1 | 0.02411 | AT2G32415.3 | AtRRP6L3 | Polynucleotidyl transferase, ribonuclease H fold protein with HRDC domain-containing protein;(source:Araport11) |
| *V. unguiculata* | Vigun07g056000.1 | 0.02789 | AT1G49350.1 | pseudoruridine kinase (PUKI) | pfkB-like carbohydrate kinase family protein;(source:Araport11) |
| *V. marina* | Vigma.0013s056600.01 | 1.38E-09 | AT1G32610.2 | - | hydroxyproline-rich glycoprotein family protein;(source:Araport11) |
| *V. marina* | Vigma.0033s033500.01 | 4.17E-09 | AT1G05060.1 | - | coiled-coil protein;(source:Araport11) |
| *V. marina* | Vigma.0001s032400.01 | 1.75E-08 | AT4G06599.1 | - | ubiquitin family protein;(source:Araport11) |
| *V. marina* | Vigma.1067s000400.01 | 0.00021 | - | - | - |
| *V. marina* | Vigma.0095s021200.01 | 0.00035 | - | - | - |
| *V. marina* | Vigma.1139s001000.01 | 0.02401 | AT3G23890.2 | topoisomerase II (TOPII) | topoisomerase II;(source:Araport11) |

**Table S9. Selected salinity- and drought-related genes upregulated in tolerant species.**

| Symbol | Gene Name | Description | Reference |
|---|---|---|---|
| *AATP1* | AAA-ATPASE 1 | ATPase induced by stress | Rama Devi et al., 2006 |
| *ABA1* | ABA DEFICIENT 1 | Zeaxanthin epoxidase involved in the first step of ABA producition. | Marin et al., 1996 |
| *ABA2* | ABA DEFICIENT 2 | Cytosolic short-chain dehydrogenase/reductase involved in converting xanthoxin to ABA-aldehyde during ABA biosynthesis. | González-Guzmán et al. 2002 |
| *ABA3* | ABA DEFICIENT 3 | Involved in convertig ABA-aldehyde to ABA, the last step of ABA biosynthesis. | Bittner et al., 2001 |
| *ABCG40* | ATP-BINDING CASSETTE G40 | ABA transporter | Kuromori and Shinozaki, 2010 |
| *ABF2* | ABSCISIC ACID RESPONSIVE ELEMENTS-BINDING FACTOR 2 | Leucine zipper transcription factor that binds to ABA–responsive element motif in the promoter region of ABA-inducible genes. | Uno et al., 2000 |
| *ABI1* | ABA INSENSITIVE 1 | Involved in abscisic acid (ABA) signal transduction | Leung et al., 1994 |
| *ACA4* | AUTOINHIBITED CA(2+)-ATPASE, ISOFORM 4 | Vacuole-localized calmodulin-regulated Ca(2+)-ATPase that improves salt tolerance. | Geisler et al., 2000 |
| *AIRP4* | ABA INSENSITIVE RING PROTEIN 4 | Cytosolic protein with E3 ligase activity that is involved in positive regulation of ABA responses. | Yang et al., 2015 |
| *ALDH3I1* | ALDEHYDE DEHYDROGENASE 3I1 | Aldehyde dehydrogenase induced by ABA and dehydration that oxidizes saturated aliphatic aldehydes. | Kotchoni et al., 2006 |
| *ANN1* | ANNEXIN 1 | Mediate Ca2+ signaling rensponding to osmotic/salt stress. | Huh et al., 2010 |
| *ANN2* | ANNEXIN 2 | Mediate Ca2+ signaling rensponding to osmotic/salt stress. | Lee et al., 2004 |
| *ANN3* | ANNEXIN 3 | Mediate Ca2+ signaling rensponding to osmotic/salt stress. | Liu et al., 2021 |
| *ANN8* | ANNEXIN 8 | Involved in multiple stress signaling pathways | Yadav et al., 2022 |
| *BGLU13* | BETA GLUCOSIDASE 13 | Beta glucosidase 13 | Xu et al., 2004 |
| *CCD1* | CAROTENOID CLEAVAGE DIOXYGENASE 1 | Encodes a protein with 9-cis-epoxycarotenoid dioxygenase activity, regulatign a key step of ABA production. | Qin and Zeevaart, 1999 |
| *CDPK6* | CALCIUM-DEPENDENT PROTEIN KINASE 6 | Required for MAPK-independent salt acclimation. | Mehlmer et al., 2010 |
| *CIPK9* | CBL-INTERACTING PROTEIN KINASE 9 | Similar to SOS2, involved in K+ homeostasis. | Liu et al., 2012 |
| *EBP* | ETHYLENE-RESPONSIVE ELEMENT BINDING PROTEIN | Member of the ERF (ethylene response factor) of the plant specific ERF/AP2 transcription factor family (RAP2.3). | Papdi et al., 2015 |
| *EIN2* | ETHYLENE INSENSITIVE 2 | Involved in ethylene signal transduction. | Alonso et al., 1999 |
| *ERA1* | ENHANCED RESPONSE TO ABA 1 | Beta subunit of farnesyl-trans-transferase, involved in meristem organization and ABA-mediated signal transduction. | Uno et al., 2000 |
| *ERF105* | ETHYLENE RESPONSIVE ELEMENT BINDING FACTOR 105 | Member of the ERF (ethylene response factor) of the plant specific ERF/AP2 transcription factor family (RAP2.3). | Illgen et al., 2020 |
| *ERF53* | ERF DOMAIN 53 | Drought-induced transcription factor of AP2/ERF superfamily. Regulates drought-responsive gene expressions. | Hsieh et al., 2013 |
| *ETR1* | ETHYLENE RESPONSE 1 | Ethylene receptor. | Wilson et al., 2014 |
| *FLZ10* | FCS LIKE ZINC FINGER 10 | Iinduced during energy starvation through SnRK1 signaling. | Jamsheer et al., 2018 |
| *GolS1* | GALACTINOL SYNTHASE 1 | Catalyzes formation of galactinol from UDP-galactose and myo-inositol. Invoved in tolerance to salt, chilling, and high-light stress. | Taji et al., 2002 |
| *GSTF10* | GLUTATHIONE S-TRANSFERASE PHI 10 | Early dehydration-induced glutathione S-transferases | Kiyosue et al. 1993 |
| *GSTU19* | GLUTATHIONE S-TRANSFERASE TAU 19 | Tau GST gene family induced by drought stress, oxidative stress, and high doses of auxin and cytokinin. | Bianchi 2002 |
| *HB-7* | HOMEOBOX 7 | transcriptionally regulated in an ABA-dependent manner and acts in a signal transduction pathway which mediates a drought response | Liu et al., 2007 |
| *HKT1* | HIGH-AFFINITY K+ TRANSPORTER 1 | Sodiun transporter involved in transferring sodium from xylem sap to phloem sap. | Sunarpi et al., 2005 |

**Table S9. Selected salinity- and drought-related genes upregulated in tolerant species, continued.**

| Symbol | Gene Name | Description | Reference |
|---|---|---|---|
| *HSFA2* | HEAT SHOCK TRANSCRIPTION FACTOR A2 | Heat Stress Transcription Factor (Hsf) family. Involved in response to misfolded protein accumulation in the cytosol | Nishizawa et al., 2006 |
| *HSFA4A* | HEAT SHOCK TRANSCRIPTION FACTOR A4A | Member of Heat Stress Transcription Factor family that is a substrate of the MPK3/MPK6 signaling and regulates stress responses. | Pérez-Salamó, et al., 2014 |
| *HSP81-2* | HEAT SHOCK PROTEIN 90.2 | Heat Shock Protein involved in sotamatal closure and ABA response. | Clément et al., 2011 |
| *ITN1* | INCREASED TOLERANCE TO NACL 1 | Protein with an ankyrin motif and transmembrane domains involved in salt tolerance. | Sakamoto et al., 2008 |
| *LEA4-5* | LATE EMBRYOGENESIS ABUNDANT 4-5 | Typically accumulates in response to low water availability conditions imposed during development or by the environment. | Olvera-Carrillo et al., 2010 |
| *LOX2* | LIPOXYGENASE 2 | Chloroplast lipoxygenase required for wound-induced jasmonic acid accumulation in Arabidopsis. Involved in stomatal closure. | Sun et al., 2015 |
| *MPK4* | MAP KINASE 4 | Phosphorylates heat shock factor A4A which regulates responses to combined salt and heat stresses | Andrási et al., 2019 |
| *NCL* | NA+/CA2+ EXCHANGER | Participates in the maintenance of Ca 2+ homeostasis | Wang et al., 2012 |
| *NHD1* | SODIUM:HYDROGEN ANTIPORTER 1 | Na+/H+ antiporter in chloroplast | Müller et al., 2014 |
| *NHX1* | NA+/H+ EXCHANGER 1 | Vacuolar Na+/H+ antiporter involved in ion homeostasis for salt tolerance. | Apse et al. ,1999 |
| *NHX2* | NA+/H+ EXCHANGER 2 | Vacuolar K+/H+ exchanger for K+ uptake, involved in regulating stomatal closure. | Cellier et al., 2004 |
| *NLP7* | NIN LIKE PROTEIN 7 | Modulates nitrate sensing and metabolism, also affects drought tolerance | Castillo et al., 2021 |
| *OST1* | OPEN STOMATA 1 | member of SnRK2 activated by salt and osmotic stress. Involved in stomatal closure. | Kawa et al., 2019 |
| *P5CR* | PYRROLINE-5- CARBOXYLATE (P5C) REDUCTASE | Catalyzes the final step in proline biosynthesis | Verbruggen et al., 1993 |
| *P5CS1* | DELTA1-PYRROLINE-5-CARBOXYLATE SYNTHASE 1 | Catalyzes the rate-limiting step in proline biosynthesis. | Strizhov et al., 1997 |
| *PIP1* | PLASMA MEMBRANE INTRINSIC PROTEIN 1 | Aquaporin. | Kaldenhoff et al., 1998 |
| *PIP2* | PLASMA MEMBRANE INTRINSIC PROTEIN 2 | Aquaporin. | Xiaojuan et al., 2011 |
| *RAP2.1* | RELATED TO AP2 1 | DREB subfamily A-5 of ERF/AP2 transcription factor family | Dong and Liu, 2010 |
| *RCD1* | RADICAL-INDUCED CELL DEATH1 | Promotes cell death when superoxide is present. | Teotia and Lamb, 2009 |
| *RD19* | RESPONSIVE TO DEHYDRATION 19 | Similar to cysteine proteinases, induced by desiccation but not abscisic acid. | Koizumi et al., 1993 |
| *RD21A* | RESPONSIVE TO DEHYDRATION 21A | Cysteine proteinase precursor-like protein/ dehydration stress-responsive gene | Koizumi et al., 1993 |
| *RD26* | RESPONSIVE TO DEHYDRATION 26 | NAC transcription factor induced by desiccation. Activates ABA-mediated dehydration response. | Takasaki et al., 2015 |
| *RGLG2* | RING DOMAIN LIGASE2 | RING domain ubiquitin E3 ligase that negatively regulates drought stress response | Cheng et al., 2011 |
| *SAT32* | SALT-TOLERANCE 32 | Involved in salt tolerance. Loss of function cause hypersensitivity to salt stress. | Park et al., 2009 |
| *SDIR1* | SALT- AND DROUGHT-INDUCED RING FINGER1 | Positive regulator of ABA signaling | Zhang et al., 2007 |
| *SIP3* | SOS3 INTERACTING PROTEIN 3 | Required for development and salt tolerance | Tripathi et al., 2009 |
| *SnRK2.4* | SUCROSE NONFERMENTING 1-RELATED PROTEIN KINASE 2-4 | Calcium/calmodulin-dependent protein kinase subfamily activated by salt and osmotic stress. | Boudsocq et al., 2004 |
| *SOS1* | SALT OVERLY SENSITIVE 1 | Functions in the extrusion of toxic Na+ from cells and is essential for plant salt tolerance | Shi et al., 2000 |
| *SOS2* | SALT OVERLY SENSITIVE 2 | Regulates SOS1 activity via phosphorylation. | Liu et al., 2000 |
| *STZ* | SALT TOLERANCE ZINC FINGER | Zinc-finger protein required for Li+ and Na+ efflux in yeast. | Lippuner et al., 1996 |
| *TEJ* | POLY(ADP-RIBOSE) GLYCOHYDROLASE 1 | Plays a role in abiotic stress responses and DNA repair | Song et al., 2015 |
